# Detecting DNA of novel fungal pathogens using ResNets and a curated fungi-hosts data collection

**DOI:** 10.1101/2021.11.30.470625

**Authors:** Jakub M. Bartoszewicz, Ferdous Nasri, Melania Nowicka, Bernhard Y. Renard

## Abstract

**Background:** Emerging pathogens are a growing threat, but large data collections and approaches for predicting the risk associated with novel agents are limited to bacteria and viruses. Pathogenic fungi, which also pose a constant threat to public health, remain understudied. Relevant data remains comparatively scarce and scattered among many different sources, hindering the development of sequencing-based detection workflows for novel fungal pathogens. No prediction method working for agents across all three groups is available, even though the cause of an infection is often difficult to identify from symptoms alone.

**Results:** We present a curated collection of fungal host range data, comprising records on human, animal and plant pathogens, as well as other plant-associated fungi, linked to publicly available genomes. We show that it can be used to predict the pathogenic potential of novel fungal species directly from DNA sequences with either sequence homology or deep learning. We develop learned, numerical representations of the collected genomes and visualize the landscape of fungal pathogenicity. Finally, we train multi-class models predicting if next-generation sequencing reads originate from novel fungal, bacterial or viral threats.

**Conclusions:** The neural networks trained using our data collection enable accurate detection of novel fungal pathogens. A curated set of over 1,400 genomes with host and pathogenicity metadata supports training of machine learning models and sequence comparison, not limited to the pathogen detection task.

**Availability:** The data, models and code are hosted at https://zenodo.org/record/5846345, https://zenodo.org/record/5711877, and https://gitlab.com/dacs-hpi/deepac.

## 1 Introduction

Many species of fungi are dangerous plant, animal, or human pathogens. Importantly, even usually harmless opportunists can be deadly in susceptible populations. For example, *Candida albicans* causes common, relatively benign infections like thrush and vulvovaginal candidosis, affecting up to 75% of women at least once in their lifetime and often re-occurring multiple times (Sobel, 2007). It is also frequently found in healthy humans without leading to any disease, and has been reported to be capable of stable colonization (Raimondi *et al*., 2019). However, invasive *Candida* infections, especially bloodstream infections, can reach mortality rates of up to 75%, rivaling those of bacterial and viral sepsis (Brown *et al*., 2012). A related species, *Candida auris*, has been first recognized in a human patient in 2009 (Satoh *et al*., 2009) and quickly became one of the most urgent threats among the drug-resistant pathogens (CDC, 2019), reaching mortality rates of up to 60% (Spivak and Hanson, 2018). It might have originally been a plant saprophyte which has adapted to avian, and then also mammalian hosts, possibly prompted by climate change (Casadevall *et al*., 2019). Strikingly, it seems to have emerged in three different clonal populations on three continents at the same time, for reasons that currently remain unexplained (Lockhart *et al*., 2017).

Even though fungal infections are estimated to kill 1.6 million people a year, they remain understudied and underreported (Chowdhary *et al*., 2016; Huseyin *et al*., 2017; noa, 2017). Estimates suggest that between 1.5 million (Hawksworth, 2001) and 5.1 million (Blackwell, 2011), or even 6 million (Taylor *et al*., 2014) different species of fungi exist, but only a small fraction of them has been sequenced. This poses a major challenge especially for pathogen detection workflows based on next-generation sequencing (NGS). Standard methods are based on recognition of known taxonomic units by homology detection, using either sequence alignment (Li and Durbin, 2010; Langmead and Salzberg, 2012; Li, 2018; Altschul *et al*., 1990; Camacho *et al*., 2009; Hong *et al*., 2014; Naccache *et al*., 2014; Ahn *et al*., 2015; Andrusch *et al*., 2018), k-mer based approaches (Wood *et al*., 2019; Piro *et al*., 2020; Breitwieser *et al*., 2018) or combinations thereof (Piro *et al*., 2017). This in turn requires curated databases of fungal, as well as bacterial, viral, and other species labelled with information regarding the corresponding pathogenic phenotype or host information. Limited host information is available in the NCBI Genome browser (Sayers *et al*., 2021a), Database of Virulence Factors in Fungal Pathogens (Lu *et al*., 2012) and the U.S. National Fungus Collections Fungus-Host Database (Farr and Rossman, 2021). Those resources are partially complementary, and none of them encompasses all the available data. What is more, multiple literature sources describe fungal pathogens and their hosts without referring to the corresponding genomes, even if they are indeed available in databases such as GenBank (Sayers *et al*., 2021b) or FungiDB (Basenko *et al*., 2018), which store genomic data without clear-cut host annotation. The ENHanCEd Infectious Diseases Database (EID2) (Wardeh *et al*., 2015) aims to detect all ‘carrier’-’cargo’ relationships, not limited to fungi or pathogens specifically, although it does contain fungal pathogens as well. It relies on automatically mining the ‘host’ field in NCBI Taxonomy (Schoch *et al*., 2020) and finding co-occurrences of species names in articles indexed by PubMed (Sayers *et al*., 2021a), providing links to the associated nucleotide sequences. This method is efficient and scalable, but automated processing based on a concise set of simplifying assumptions may sometimes lead to spurious results. Many ‘cargo’ and ‘carrier’ species can be mentioned in the same paper even though one is not really a host of the other. This is often the case in literature reviews, taxonomy updates, and holds also for this work. The ‘host’ field in a database as large as NCBI Taxonomy may also contain outdated, inaccurate or incomplete information. For example, *Pneumocystis jirovecii*, the causative agent of deadly pneumocystis pneumonia, was previously called *P. carinii*. While the latter name is now reserved for a species infecting exclusively rats and not humans (Stringer *et al*., 2002), records in Taxonomy (and, possibly by consequence, EID2) still list humans as the hosts of *P. carinii* at the time of writing. What is more, many sequences included in EID2 are not genome assemblies, but single genes, which are not enough for open-view fungal pathogen detection based on shotgun sequencing. For this and similar applications, a new resource is needed.

We compiled a collection of metadata on a comprehensive selection of fungal species, annotated according to their reported host groups and pathogenicity. We store the metadata in a flat-file database and link them to the corresponding representative (as defined in GenBank) or reference genomes, if available. To showcase the possible applications of the database, we model a scenario of novel fungal pathogen detection. While to our knowledge, this is a first systematic evaluation of feasibility of this task, we note that it mirrors similar problems in bacterial and viral genomics (Deneke *et al*., 2017; Mock *et al*., 2020; Zhang *et al*., 2019; Barash *et al*., 2018; Guo *et al*., 2021; Gałan *et al*., 2019; Wardeh *et al*., 2021; Brierley and Fowler, 2021; Bergner *et al*., 2021; Tang *et al*., 2015; Bartoszewicz *et al*., 2020, 2021b). We expect new agents to emerge due to environmental changes, host-switching events and growing human exposition to the unexplored diversity of potentially harmful fungi, as shown by the example of *C. auris*. Further, advances in engineering of fungal genomes (Richardson *et al*., 2017; Burgess, 2017; Dai *et al*., 2020; Szymanski and Calvert, 2018; Luo *et al*., 2018; Amores *et al*., 2016; Martins-Santana *et al*., 2018) could lead to new risks, and screening of synthetic sequences relies on of methods developed originally for pathogen detection (Diggans and Leproust, 2019; Balaji *et al*., 2021). Therefore, we evaluate if detecting homology between previously unseen species and their known relatives accurately predicts if a DNA sequence originates from a novel fungus capable of colonizing and infecting humans. BLAST (Altschul *et al*., 1990; Camacho *et al*., 2009) represents the gold standard in pathogen detection via taxonomic assignment to the closest relative. Although read mappers or k-mer based taxonomic classifiers are more computationally efficient on large NGS datasets (Breitwieser *et al*., 2017; Ye *et al*., 2019; Alser *et al*., 2021), BLAST has been shown to be more accurate in similar tasks of detecting novel bacterial and viral pathogens (Deneke *et al*., 2017; Bartoszewicz *et al*., 2021b). However, convolutional neural networks of the DeePaC package have been proven to outperform BLAST in both those scenarios (Bartoszewicz *et al*., 2020, 2021b) for both isolated NGS reads and full genomes, and a recently presented variant of residual neural networks (ResNets) outperforms all alternatives on short NGS reads and their fragments (Bartoszewicz *et al*., 2021a). We trained similar ResNets to predict if a novel DNA sequence originates from a human-infecting fungus. To visualize the dataset, we developed trained numerical representations of all genomes in the database.

## 2 Methods

### 2.1 Data description

We collected metadata on species infecting humans, animals or plants, supplemented with information on other plant-associated species. To this end, we integrated multiple literature and database sources (see Supp. Information for citations), relying on manual curation, but also including the automatically extracted data for future reference. Table S1 summarizes the data we collected for each species. We describe the curation procedure in detail in Supp. Note 1. The database contains 14,555 records in total. Here, we will focus on what we will call the core database, comprising metadata on genomes of 954 manually confirmed pathogens (including 332 species reported to cause disease in humans), available on October 9, 2021. This forms a collection of species most relevant to the pathogen detection task, belonging to 6 phyla, 37 classes, 82 orders and 182 families. A ‘temporal benchmark’ subset contains 15 further pathogens (including one infecting humans), collected in a database update on January 2, 2022. We also include records on 486 plant-associated fungi. The supplementary part of the database contains information on 481 putatively labelled genomes, 1,147 unlabelled species with available genomes, 8 synonyms (with 6 alternative genomes) derived from the Atlas of Clinical Fungi (de Hoog *et al*., 2020), 885 labelled species without genomes (including 284 species without TaxIDs) and 10,579 putatively labelled species without genomes (including 9 without TaxIDs). This subset will enable easy updating of the database in the future, as more genomes of already labelled species are sequenced. It also serves as a record of all screened genomes and species to ensure reproducibility and facilitate future extensions (e.g. adding new data or sources of evidence). Figure S1 presents the numbers of genomes with manually confirmed labels and genomes for which putative labels could be found in EID2 (Wardeh *et al*., 2015).

### 2.2 Training, validation and test sets

While we envision a wide range of possible applications of the database, we present example usecases allowing to take advantage of the wealth of collected data – detection of novel fungal pathogens from NGS data. The core of the database contains 332 genomes of human pathogens (including opportunists), forming the positive class. The negative class comprises 622 species not reported to infect humans; this includes 565 plant pathogens and 58 non-human animal pathogens. To evaluate the performance of the selected pathogenic potential prediction methods, we divided the corresponding genomes into non-overlapping training, validation and test sets. In this setup, the training set is used as a reference database for the methods based on sequence homology and to train the neural networks, while the performance metrics are calculated on the held-out test set. The validation set is used for hyperparameter tuning and to select the best training epoch. While evaluating performance on genuinely unknown species is by definition impossible, we effectively model the ‘novel species’ scenario by testing on a wide range of sequences removed from the database, including selected important species (Satoh *et al*., 2009; Brown *et al*., 2012; Dean *et al*., 2012; Skamnioti and Gurr, 2009; Scheele *et al*., 2019). We simulated low- and high-coverage read sets using a protocol outlined in Supp. Note 2 (Holtgrewe, 2010). We considered two methods of balancing the number of samples between taxa: setting the read number per species to be proportional to the respective genome’s length (‘linear-size’) or its logarithm (‘logarithmic-size’).

### 2.3 Phenotype prediction and genome representations

Next, we evaluated the feasibility of pathogenic potential prediction for novel fungal species. We used a ResNet architecture implemented in the DeePaC package, previously shown to outperform alternatives based on deep learning, traditional machine learning and sequence homology in the context of novel bacteria and viruses (Bartoszewicz *et al*., 2021a). We also adapted a BLAST-based pipeline used by Bartoszewicz *et al*. (2020) for benchmarking. Details of the pipeline, the ResNet architecture and the hyperparameter tuning procedure are described in Supp. Note 3.

To visualize the structure of the dataset as learned by the trained classifier, we developed numerical representations for the collected genomes. This poses a challenge, as the networks are trained on reads rather than full genomes. However, we observe that final outputs of the network can be averaged over all reads originating from a single genome to generate a prediction for the genome in question (Bartoszewicz *et al*., 2020). Analogously, we can average the activations of the intermediate layers to construct vector representations for whole genomes based on the corresponding reads. Note that averaging the activations of the penultimate layer is approximately equivalent to using a full genome as input (assuming full coverage), as our architecture uses global average pooling just before the output layer. Classifier outputs based on average activations indeed approximate classifier outputs averaged over all reads originating from a given species (see Supp. Note 4 and Figure S2-S3). Hence, we extracted penultimate activations for all simulated reads in the low-coverage, ‘linear-size’ training, validation and test sets. We then used the averaged activation vectors for each species to map the distances between them as learned by our networks. We used UMAP (McInnes *et al*., 2020) to visualize the dataset (Supp. Note 6).

### 2.4 Multi-class evaluation

Finally, we investigated an application requiring merging the ‘positive’ subset of our database with previously available resources for pathogenic potential prediction in bacteria and viruses. We aimed to integrate the separate classifiers for fungi, bacteria and viruses into a single, multi-class model capable of predicting whether unassembled NGS reads originate from (possibly novel) pathogens present in a human-derived sample. To this end, we extended the DeePaC package adding the multi-class classification functionality. The resulting architectures differ from the previously described ResNets (Bartoszewicz *et al*., 2021a) only by the output layer, which has as many units as the number of considered classes and uses a softmax activation (see Supp. Note 3). In practice, only human-hosted fungi are expected to be found in clinical samples. In this context, a slightly constrained view is admissible: we assume that only human-pathogenic fungi, human-hosted bacteria (pathogenic or commensal), human viruses and non-human viruses (mainly bacteriophages) will be present in the sample. Further, bacteriophage sequences tend to be very similar to the sequences of their bacterial hosts (Zielezinski *et al*., 2021b,a) and difficult to differentiate, but both commensal bacteria and non-human viruses can be viewed here as a joint ‘negative’ (i.e. harmless) class. Hence, learning a precise decision boundary between them can be omitted. Human reads can be ignored, as they can be relatively easily filtered out with traditional methods based on read mapping or k-mers (Loka *et al*., 2018; Ahmed *et al*., 2021; Wood *et al*., 2019). Therefore, we fused the previously published datasets used in DeePaC (Bartoszewicz *et al*., 2021a) for bacteria (pathogens vs. commensals) and viruses (human vs. non-human) with the ‘positive’ (human-pathogenic) class of our database (Supp. Notes 2-3).

The final result is a dataset divided into 4 classes: nonpathogenic bacteria and non-human viruses, bacterial pathogens, human-infecting viruses, and human-pathogenic fungi, in either the ‘linear-size’ or the ‘logarithmic-size’ variant. Note that even in this case, the ‘negative’ part of the presented database is useful, allowing us to constrain our view to a curated set of clinically relevant fungi only. Using this dataset, we trained two models including all four classes (using the ‘linear-size’ or the ‘logarithmic-size’ variant of the fungal training set). We further evaluated the one resulting in higher validation accuracy and a simple ensemble averaging the predictions of both models. Then, we trained a 3-class model including only the bacterial and viral classes. This allows us to measure the ‘difficulty’ of integrating the fungal dataset with the others within a single network in terms of resulting differences in prediction accuracy on the original DeePaC datasets. By comparing the performance of our models to the performance of the original binary classifiers (Bartoszewicz *et al*., 2021a), we can disentangle the ‘difficulty’ of adding the fungal class from the ‘difficulty’ of integrating the bacterial and viral classes, and assess how much performance is ‘lost’ by using a more open-view classifier. Note that in the case of the purely viral dataset, spurious assignments to the bacterial pathogens class may be treated as detection of bacteriophages infecting the bacterial species of this class, and hence reassigned into predictions for the non-pathogen class by adding the predicted probabilities for both classes. This effectively merges the non-pathogen and bacterial pathogen classes at test time when appropriate, but still keeps the possibility to use the trained networks in a fully open-view setting (with all classes) without the need for retraining. We performed an additional comparison to BLAST with a pre-selected training database (bacterial for bacteria (Bartoszewicz *et al*., 2020), viral for viruses (Bartoszewicz *et al*., 2021b)). This resulted in an estimated upper bound on the performance of non-machine learning approaches on those datasets (as extending the training database with irrelevant reference genomes can only lower BLAST’s performance). Finally, we evaluated the neural networks on the full dataset of all four classes and a real *C. auris* sequencing run. We also analyzed the latter with STAT (Katz *et al*., 2021), used by the SRA database (Supp. Note 7).

## 3 Results

### 3.1 Fungal pathogenic potential prediction

The best network, trained on the high-coverage, ‘logarithmic-size’ dataset without dropout, required 8 days of training on four Tesla V100 GPUs and was selected for further evaluation. Proper retuning of the classification threshold for species-level predictions appears to be a necessary step for an independent, viral dataset (Table S2), so we also retuned the threshold (0.46 instead of the default 0.5) for the respective fungal ResNet setup. Overall, prediction accuracy for reads and read pairs is suboptimal for both BLAST and the ResNet, probably reflecting the extreme difficulty of the task (Supp. Note 5, Table S3). The error estimates based on the held-out test dataset are consistent with the results of the temporal benchmark (Table S4). Balanced accuracy is much higher on full genomes (88.4– 90.3), suggesting that the main performance bottleneck is the total amount of information (total sequence length) available as input. This is consistent with previous observations (Deneke *et al*., 2017; Bartoszewicz *et al*., 2020, 2021b), although more drastic than in the case of bacteria or viruses. All approaches correctly classify the single human pathogen in the temporal benchmark dataset, but the ResNet achieves higher specificity (92.9) than read-based (78.6) and contig-based BLAST (85.7). Despite the low coverage, the test set seems to be indeed representative. The genome-wide predictions are equally accurate if all test reads are used, and when only a half of the dataset (corresponding to either the first or the second mate) is analyzed. Therefore, computations for single-species samples can be sped up by considering first mates only, as they are enough to deliver an accurate prediction. Strikingly, this is the case even though they correspond to a mean coverage below 0.08. As a result, the read-based ResNet yields only slightly less accurate predictions than contig-based BLAST, but requires 700-fold less time if a GPU is used (Supp. Note 5, Table S3).

### 3.2 The landscape of fungal pathogenicity

Good overall accuracy of the ResNet is reflected in the visualisation of learned genome representations for the entirety of the core database. Figure 1 and Figures S4–S10 presents UMAP embeddings of the extracted representations for all labelled genomes, i.e. a sum of the training, validation and test datasets. Although some noise is present, the positive and the negative class are mostly separated. Several clusters of human pathogens and non-human pathogens are present. The ResNet correctly recovers most of the labels, including many of the ‘positive’ members of the otherwise ‘negative’ clusters (Figure S5). To measure this, we visually identified 14 clusters, which could be easily retrieved automatically using single-linkage agglomerative clustering (Figure S10). Cluster purity for the whole dataset was high (0.90). We also measured it for the members of the large, mixed clusters 4 (pink in Figure S10) and 6 (red in Figure S10), which achieved purity of 0.85 and 0.88, respectively. Classification errors seem to originate from an interpolation based on neighbouring data points – within clusters, the predicted labels are more homogeneous than the ground truth annotations. This is expected, as the clusters represent similarity in the space of learned representations. The network should in general assign similar labels to inputs similar in this space. In contrast, BLAST works analogously to a k-nearest neighbours classifier in the input sequence space (finding the single closest match for each query). The ResNet, interpolating between multiple data points, may be less efficient in modelling situations where a small set of ‘negative’ data points is embedded within a larger ‘positive’ cluster of similar species, or vice versa. This hypothesis is supported by the visualization of BLAST-predicted labels in the learned representation space (Figure S6). BLAST recovers mixed, contrasting labels within cluster more accurately, and its errors seem to be more evenly distributed across the space. At the same time, its slightly lower sensitivity is especially visible within the diverse *Sordariomycetes* class placed in the rightmost cluster. The clusters themselves are noticeably related to the taxonomic units represented in the database, although this is importantly not a simple one-to-one mapping (Supp. Note 6, Figures S7–S10).

**Fig. 1:**
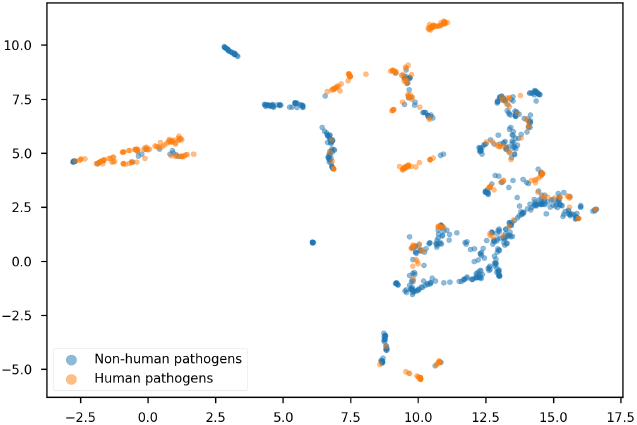
UMAP embeddings of the learned genome representations for the core database, enlarged in Figure S4. Each point represents a genome of a single species, coloured by its ground truth label. The learned representations offer a way of visualizing the core database along relevant labels for each genome. The ResNet correctly classifies most of the genomes (Figure S5). The clusters are related, but not fully reducible to the taxonomic classification of the analysed species (Supp. Note 6, Figures S7–S10).

### 3.3 Multi-class models

For the final evaluation of our database, we aimed to develop a model capable of classifying NGS reads originating from novel viruses, bacterial, and fungal species into appropriate pathogen and non-pathogen classes. We trained the multi-class ResNets on data including four classes (human-pathogenic fungi, bacterial pathogens, human viruses and non-pathogens). The network trained on the dataset containing the ‘logarithmic-size’ version of the fungal positive class achieved slightly better validation accuracy and was selected for further evaluation, but the difference was small (<0.5%). We then evaluated a simple ensemble of both 4-class ResNets. First, we used the DeePaC datasets consisting of bacteria and viruses to compare the 4-class models to a classifier including the three non-fungal classes only, as well as the binary ResNets (Bartoszewicz *et al*., 2021a) and BLAST. This procedure allows us to a) measure the effect of integrating the fungal dataset with the bacterial and viral data in one task, and b) disentangle the effects of adding the fungal data from the effects of merging the bacterial and viral datasets. We expected the fungal sequences to be relatively easy to differentiate from the others, but whether the ResNet architecture would be expressive enough to accurately represent all those diverse sequences was unclear. As shown in Table S5, integrating the fungal dataset with three bacterial and viral classes indeed does not negatively influence the prediction accuracy. BLAST, using an appropriate reference database and representing the estimated upper bound on performance of homology-based approaches, is still outperformed by a significant margin. The fungal dataset can be integrated with the other classes without causing any significant performance hits on the full, multi-class dataset as well (Table S6). Consistently with the results presented in Table S5, performance is lower on the non-pathogen class, since many bacteriophage reads can be confused with pathogenic bacteria. While this issue requires further research, we expect future solutions to remain compatible with our database. The 4-class ensemble achieves the most balanced performance on non-pathogen data, the best recall on fungal reads and is also the most accurate overall, cutting the average error rate by over 40% compared to BLAST (Table 1). As expected, distinguishing human-pathogenic fungi from the other classes is easier than predicting fungal hosts, so the performance of both BLAST and the ResNet is higher than in Table S3. This holds also for real data (Supp. Note 7, Table 2). If the correct reference genome is missing, STAT is unable to classify most of the reads (Table 2, Table S7). This is true even though genomes of related species are present in the database. The ResNets perform markedly better than BLAST. Although the simulated test sets are more representative and effectively model mock metagenomic samples, this case study shows that our methods accurately classify real data as well. The database enabled us to accurately predict whether NGS reads originate from novel pathogens.

**Table 1.** Performance on the multi-class dataset, read pairs.

The 4-class classifier includes the fungi class along the three viral and bacterial classes included in the 3-class classifier.
|  |  | Acc. | F1 | Prec. | Rec. | AUPR |
| --- | --- | --- | --- | --- | --- | --- |
| All classes | 4-class ens. (ours) | **87.6** | **87.7** | 87.7 | **87.6** | **93.4** |
|  | BLAST | 78.3 | 84.0 | **90.6** | 78.3 | - |
| Non-pathogens | 4-class ens. (ours) | **77.4** | **78.7** | **80.1** | **77.4** | **86.7** |
|  | BLAST | 66.5 | 71.6 | 77.5 | 66.5 | - |
| Path. bacteria | 4-class ens. (ours) | **87.2** | 85.1 | 83.2 | **87.2** | **90.4** |
|  | BLAST | 83.8 | **87.5** | **91.6** | 83.8 | - |
| Human viruses | 4-class ens. (ours) | **90.9** | **93.7** | 96.7 | **90.9** | **98.4** |
|  | BLAST | 78.9 | 87.9 | **99.2** | 78.9 | - |
| Fungi | 4-class ens. (ours) | **95.0** | **92.9** | 90.9 | **95.0** | **97.9** |
|  | BLAST | 84.1 | 88.9 | **94.2** | 84.1 | - |

**Table 2.**
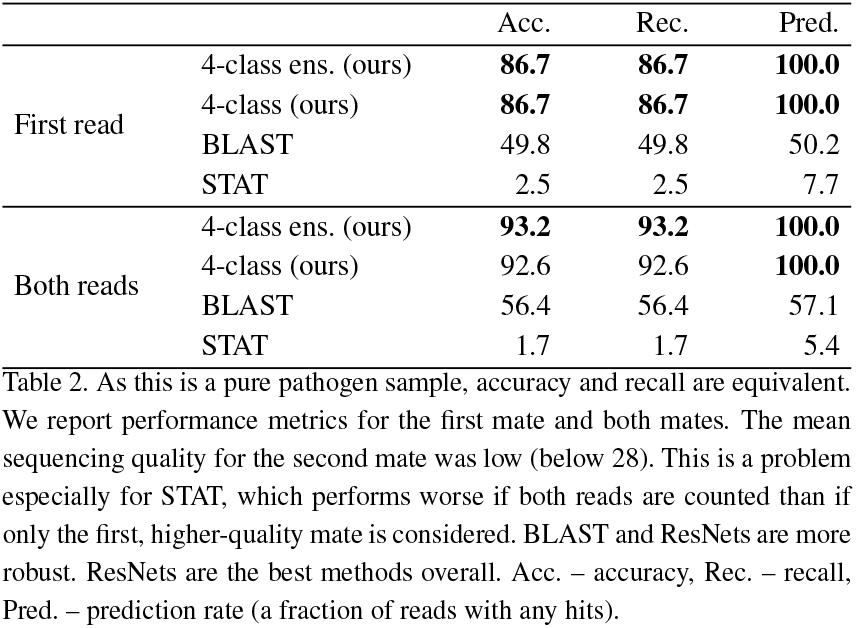
C. auris sequencing run, SRA accession SRR17577041.

As this is a pure pathogen sample, accuracy and recall are equivalent.
|  |  | Acc. | Rec. | Pred. |
| --- | --- | --- | --- | --- |
| First read | 4-class ens. (ours) | **86.7** | **86.7** | **100.0** |
|  | 4-class (ours) | **86.7** | **86.7** | **100.0** |
|  | BLAST | 49.8 | 49.8 | 50.2 |
|  | STAT | 2.5 | 2.5 | 7.7 |
| Both reads | 4-class ens. (ours) | **93.2** | **93.2** | **100.0** |
|  | 4-class (ours) | 92.6 | 92.6 | **100.0** |
|  | BLAST | 56.4 | 56.4 | 57.1 |
|  | STAT | 1.7 | 1.7 | 5.4 |

## 4 Discussion

Fungal pathogens have been under-studied compared to human-infecting bacteria and viruses, leading to repeated calls for more research in this area (Huseyin *et al*., 2017; noa, 2017). What is more, a large part of the research effort has been focused on plant pathogens due to their agricultural significance. A subset of them could in principle also have an unreported or undetected ability to infect a human host. An analogous problem applies also to incomplete data regarding pathogenicity towards non-human animals or plants. For this reason, we do not claim that species not listed as potential pathogens are indeed non-pathogens. In our database, we include confirmed labels alongside appropriate sources; in case of lack of evidence, we treat the respective label as missing. It is therefore possible that some of the fungi currently labelled as ‘non-human pathogens’ would have to be reclassified as the state of science evolves. This may be especially important as it has been suggested that the ongoing climate change will lead to more frequent host-switching events, including expansion of host-range to mammals, which are usually relatively resistant to fungal infections (Garcia-Solache and Casadevall, 2010). Even though the very goal of the presented classifiers is to generalize to newly emerging species, large, comprehensive datasets are crucial – often more important than the actual analysis method used. This has been shown before for metagenomic data (Piro *et al*., 2020) and likely applies to the tasks analyzed here as well. Therefore, extending the database to include more species, as more genomes are sequenced in the future, could facilitate the downstream tasks. To support future extensions, we include all considered species in the database – even those without assigned TaxIDs, genomes, or labels (in the case of screened GenBank genomes). This broadens the scope of the data from 1,455 labelled genomes to over 14,500 records, enabling easy labelling of newly published genomes and minimizing the workload needed for addition of new, non-redundant records. It is also possible to link the species TaxIDs to taxa below the species level. However, it should be kept in mind that the fungal taxonomy is in constant flux – taxa previously considered variants of a single species may be reclassified into separate species in the future. While automatically curated databases like EID2 (Wardeh *et al*., 2015) are relatively easy to update and scale, we note that they may be prone to errors introduced by the automated protocol used. Manual curation is not fully error-free either, but we see it as a necessary step to maximize the quality of the collected labels. Both approaches are complementary and may be best suited for different use-cases.

We show that both BLAST and ResNet can accurately predict if a fungus is a human pathogen based on its genome. The read-level performance is admittedly low for predicting a fungal host, but the trained representations allowed us to visualize the taxonomic diversity of the database along its phenotypic landscape. As expected, the apparent fungal host-range signal seems to be related to, but not fully reducible to the fungal taxonomy. Most importantly, multi-class networks detecting fungal, bacterial or viral pathogens noticeably outperform the homology-based approach. Different extraction protocols can affect the relative yield of bacterial and fungal DNA (Fiedorová *et al*., 2019), but the methods investigated here process one sequence at a time, so are not affected by this kind of bias. Further work could extend the presented multi-class setup to Nanopore reads, as shown for bacterial and viral models (Bartoszewicz *et al*., 2021a), enabling selective sequencing of mixed-pathogen samples.

Full genomes can be represented by aggregating representations of reads originating from each genome. In addition to that, we observe that coverage as low as 0.08 is enough to correctly classify a species. Taken together, those two facts warrant a view of a species genome as a distribution generating subsequences (i.e. reads) originating from it; such a distribution can also be considered in an abstract representation space (Supp. Note 4). This concept is very similar to that of a k-mer spectrum, where an *empirical* distribution of k-mers is used as a signature of a longer sequence to enable alignment-free comparisons (Zielezinski *et al*., 2017), including being used as input features for machine learning approaches as in Deneke *et al*. (2017). However, k-mer spectra operate in the sequence space only. Classifiers based on aggregated *representations* are approximately equivalent to classifiers based on aggregated *predictions*, although this relation is modulated by the standard deviation of the respective, genome-specific distribution. A somewhat related effect was reported in the context of competing design choices for neural networks equivariant to DNA reverse-complementarity – models averaging the predictions for both DNA strands were found to be approximately equivalent to models applying a sigmoid transformation to an average of logits (Zhou *et al*., 2021). The probabilistic view of genome representations presented here deserves deeper investigation; this could potentially lead to a development of useful embeddings also for whole, multi-species samples.

Although we focus on using the collected data in a pathogenic potential prediction task, the database itself can find future applications beyond this particular problem. Genomes collected here can be a valuable resource for functional and comparative genomics of fungi. For example, fungal genomes could be scanned for regions associated with their ability to colonize and infect humans, as shown previously for bacteria and viruses (Bartoszewicz *et al*., 2021b). On the other hand, the multitude of genomic features present in fungal genomes, often including intron features and regions without obvious functional annotation, renders the validation of such an approach a challenging project on its own. This could be perhaps facilitated by focusing exclusively on coding regions, which should in principle carry a stronger phenotype-related signal, at the risk of omitting potentially relevant, non-coding (e.g. regulatory) elements. As a source of curated labels, the dataset could also support application of proteomics to fungal pathogen research. Computational metaproteomics and proteogenomics approaches enable analysis of microbial communities based on mass spectrometry data and can be co-opted for pathogen detection workflows independent of DNA sequencing (Renard *et al*., 2012; Schiebenhoefer *et al*., 2019, 2020).

In conclusion, we compiled a comprehensive database of fungal species linked to their host group (human, non-human animal or plant), evidence for their pathogenicity, and publicly available genomes. To highlight the potential uses of the dataset, we benchmark two most promising approaches to novel fungal pathogen detection: a deep neural network capable of fast inference directly from DNA sequences, and the gold standard in homology-based pathogen identification – BLAST. The database, hosted at https://zenodo.org/record/5846345, can be reused for future research on fungal pathogenicity. The models, read sets and code are available at https://zenodo.org/record/5711877, https://zenodo.org/record/5846397, and https://github.com/dacs-hpi/deepac.

## Acknowledgements

We thank the NCBI help desk for assistance and helpful suggestions, as well as Katharina Baum (HPI) for valuable discussions and comments.

## Supplementary Information

### Supplementary Note 1: Database construction

First, we accessed the Database of Virulence Factors in Fungal Pathogens (DFVF) [Lu et al., 2012] on October 9, 2021. The database contains records on virulence factors of a wide selection of fungal species together with their NCBI TaxIDs and information on their hosts and the disease caused. It also includes separate parts for animal and plant pathogens. We searched the records for all proteins in the database; if the “Disease” or “Disease host” fields contained the word “human”, we added the corresponding species to the list of human pathogens. We added remaining species from the animal part of the database to the list of animal pathogens. Finally, we put all species mentioned in the plant part of the database on the list of plant pathogens, irrespective of whether humans were also mentioned as a possible host. For all those species, we also extracted the TaxID. As some of those TaxIDs corresponded to taxa below the species level, we linked them to their species-level ancestors in the NCBI Taxonomy tree. This resulted in a list of unique, labelled species names and species-level TaxIDs. Note that as DFVF was built using manual curation of text-mining results, we treat the obtained records as manually curated as well.

Further, we searched the Taxonomy database [Schoch et al., 2020] using the query ‘“host human”[Properties] AND Fungi [orgn]’, obtaining a list of species with a putative human host and their TaxIDs. We then used a comprehensive selection of literature reviews and comparative studies [Taylor et al., 2001, Chowdhary et al., 2014, Bombassaro et al., 2020, Brunner-Mendoza et al., 2019, Franzen and Müller, 2001, Colombo et al., 2011, Han and Weiss, 2017, Hu et al., 2015, Jančič et al., 2015, Kjærbølling et al., 2018, Köhler et al., 2015, Kwon-Chung et al., 2017, Paulussen et al., 2017, Prakash et al., 2017, Dellière et al., 2020, Teixeira et al., 2017, Seyedmousavi et al., 2018] to collect additional evidence and extend the collected list of human pathogens. We manually extracted the species names and searched the NCBI Taxonomy (including synonyms) to link them to their names as reported in Taxonomy and the respective species-level TaxIDs. In both cases (searching with extracted TaxIDs or species names), we handled the possibility of multiple, ambiguous matches by preferring exact name matches to any of the synonymous names recorded in Taxonomy. If this was not possible (e.g. because of additional annotations in the “name” field in Taxonomy), we considered matches where the first two words exactly match the query name, and varieties of a query species (i.e. matches containing the abbreviation ‘var.’). If no exact match was found, we selected the first record containing the query species name. We ignored species hybrids and *formae speciales*, excluded the organism names explicitly expressing taxonomic ambiguity (marked with ‘sp.’, ‘cf.’ or ‘aff.’), and filtered out fungal viruses. If the species name was not mentioned in Taxonomy at all, we still included it in the database along its associated host group label. This will enable easy extension of the database in the future, as more fungal sequences become available.

Similarly, we consulted the literature on animal [Seyedmousavi et al., 2018, Smith, 2006, Bałazy et al., 2008, Becnel and Andreadis, 2014, Franzen and Müller, 2001, Cissé et al., 2021, Chandler et al., 2000, Evans et al., 2011, Gerson et al., 2008, Han and Weiss, 2017, Schulenburg and Félix, 2017, Leonhardt et al., 2018, Lovett and St. Leger, 2017, Rehner and Buckley, 2005, Seyedmousavi et al., 2015, Shang et al., 2015, Wu et al., 2021, St. Leger and Wang, 2020, Sung et al., 2007, Teixeira et al., 2017, van der Geest et al., 2000] and plant pathogens [Vacher et al., 2008, Doehlemann et al., 2017, Barbara and Clewes, 2003, Costa et al., 2021, Coutinho et al., 2017, Gurung et al., 2015, Jones and Baker, 2007, Kim et al., 2012, 2019, Leonhardt et al., 2018, Schardl et al., 2013, Stukenbrock et al., 2012] and linked the results to the respective species TaxIDs wherever possible. Further, we accessed the Fungal Databases of the Germplasm Resources Information Network (GRIN). On October 9, 2021, we searched for all records with ‘Homo sapiens’ as a host [Farr and Rossman, 2021]. We also added the species listed in the Nomenclature Fact Sheets for plant-associated fungi with quarantine importance and the Fungal Diagnostic Fact Sheets of invasive and emerging fungal pathogens [Farr and Rossman, 2021] to the list of plant-pathogenic species, and searched NCBI Taxonomy as described earlier.

We then linked the resulting lists of species TaxIDs with their representative (as defined in GenBank) or reference genomes in GenBank, accessed on October 9, 2021. Further, we selected all species with available genomes, but without a label assigned by any of the sources used. On the same day, we manually searched the GRIN Fungal Database for each of those species, allowing for synonyms. We checked the “Host, “Disease” and “Notes” fields; if the disease of a plant or animal host was clearly described, we added the query species to the appropriate list. If no disease was mentioned for a confirmed plant host, we added the species to a list of plant-associated fungi. Next, we used the EID2 database [Wardeh et al., 2015] to extract species with putative human, animal, or plant hosts. Note that those labels were collected automatically and are therefore prone to errors. Moreover, they do not consist of pathogens only, and may include commensals or symbionts. We nevertheless added them to our database for completeness, albeit noting that they represent only a putative, automatically extracted carrier-cargo relationship, rather than a manually confirmed, true pathogenic potential of a species. The same, ‘putative’ category includes human-hosted species found in NCBI Taxonomy and GRIN, unless confirmed in other literature sources as well. We then selected all species with a putative human host and manually searched the Atlas of Clinical Fungi [de Hoog et al., 2020] with their names and synonyms noted in NCBI Taxonomy, considering also additional species referred to in the search results. If the Atlas confirmed a species to be a pathogen, we added it to the appropriate category. Note that only three of the human-hosted species (all belonging to the genus *Malassezia*) were clearly described as human commensals without any reports of causing disease (and hence, we excluded them from our list of pathogens). In cases when the Atlas mentioned two names to be synonyms, even though NCBI Taxonomy listed them as two separate species, we followed the nomenclature suggested by the Atlas. We retained both records to explicitly reflect this in the database, keeping one name unlabelled and linking it to the main record with an appropriate annotation regarding the synonym TaxID and name. Finally, we linked all labelled species to their GenBank genomes, if available.

After 12 weeks, we updated the database by extracting the records on reference and representative fungal genomes present in GenBank on January 2, 2022, selecting those with accession numbers and TaxIDs not already present in our database, and filtering the organism names as described above. We then linked the 67 new genomes to the previously collected metadata (29 of those species were absent from the first version of the database), and manually searched GRIN following the same procedure as for the main part of the database. If a human host was mentioned, we added the corresponding putative label to the appropriate list. Finally, we checked all putative human host labels in the Atlas of Clinical Fungi [de Hoog et al., 2020]. The update added 27 new labelled genomes, including 12 plant-associated fungi without proven pathogenic labels. One plant pathogen and four plant-associated species were absent from the first version of the database. We used the added pathogen genomes for our temporal benchmark.

**Figure S1:**
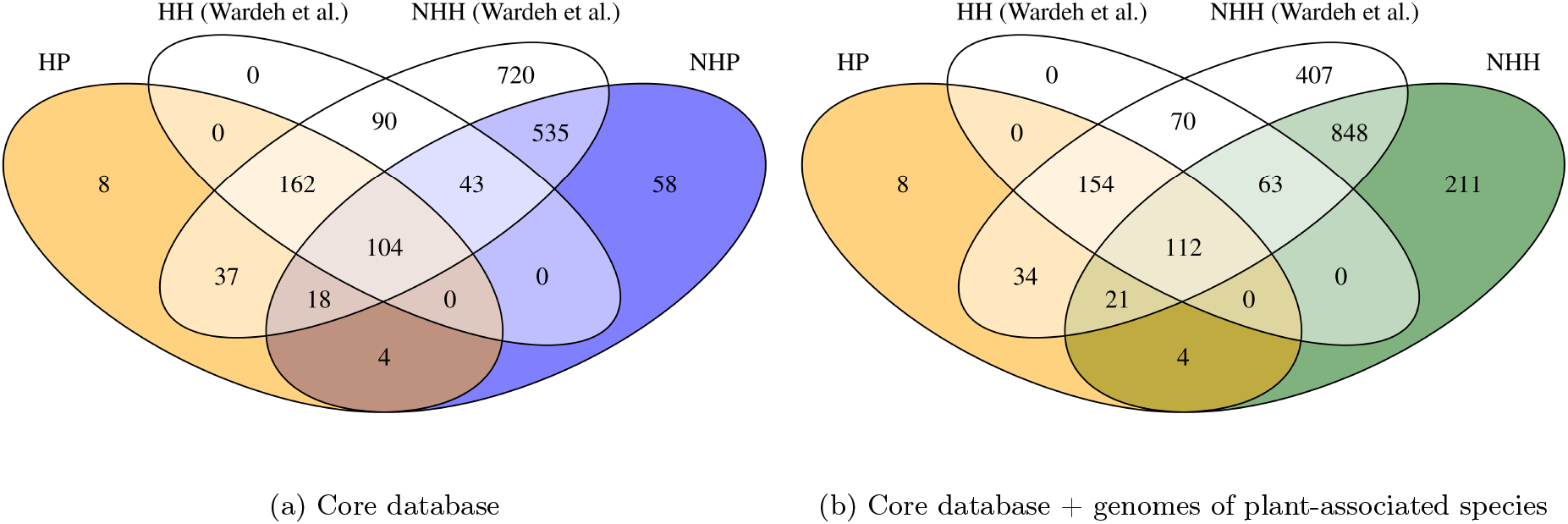
A Venn diagram of the manually confirmed labels and putative labels from EID2 [Wardeh et al., 2015] for all labelled genomes (except synonyms). HP – human pathogens (manually curated); NHP – non-human pathogens (manually curated); NHH – species with a non-human host (manually curated); HH (Wardeh et al.) – species with a putative human host (EID2); NHH (Wardeh et al.) – species with a putative non-human host (EID2).

**Table S1:**
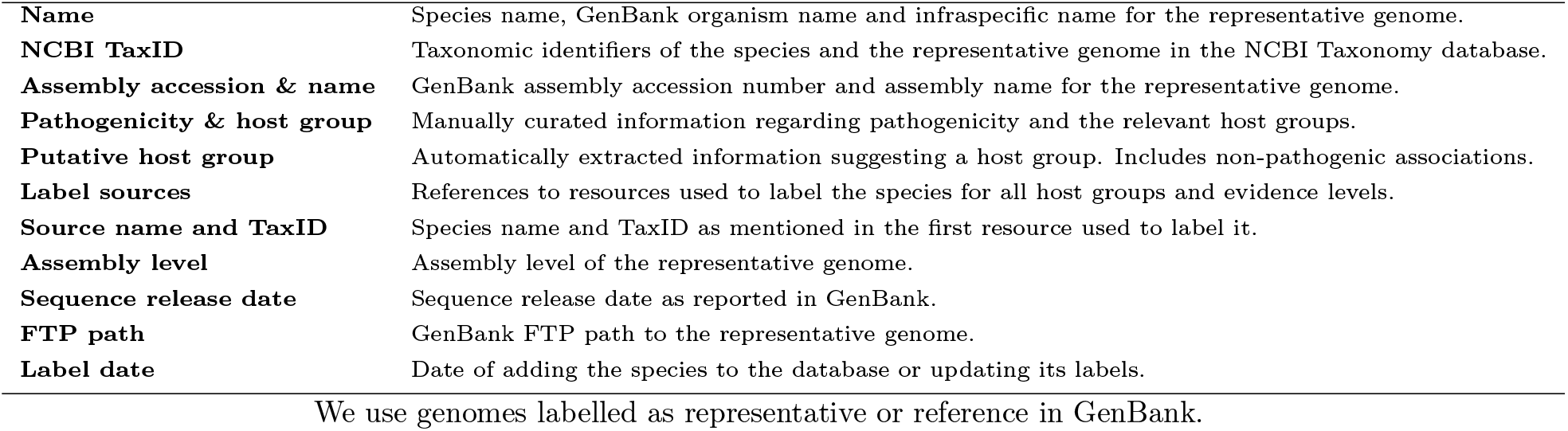
Key information on each species included in the database.

### Supplementary Note 2: Training, validation and test read set simulation

#### Fungal datasets

Pathogen ‘novelty’ is an inherently ambiguous concept. We operationalize it in taxonomic terms, measuring generalization to ‘novel’ (i.e. unseen in training) species. This is consistent with previous works (e.g. Deneke et al. [2017], Zhang et al. [2019], Bartoszewicz et al. [2020, 2021b], Barash et al. [2018], Katz et al. [2021], Wood et al. [2019]), some of which also investigated ‘novelty’ measured at other levels of taxonomy (e.g. strain). Alternatively, novelty could be expressed in terms of sequence similarity. However, designation of novel species is done by experts considering multiple criteria, including, but not limited to genome similarity. Hence, we follow the established operational definition of a novel pathogen as a novel *species*. Note that BLAST detects sequence similarity, so its performance (or lack thereof) can be interpreted as an alternative measure of ‘novelty’ of the test genomes relative to the training database.

We manually placed two clinically important pathogens – *Candida auris* (a recently emerged threat [Satoh et al., 2009, CDC, 2019]) and *Aspergillus fumigatus* (a causative agent of aspergillosis, reaching up to 95% mortality rates [Brown et al., 2012]) – in the test set. We also added two non-human pathogens: *Pyricularia oryzae* (syn. *Magnaporthe oryzae*; voted the most important fungal pathogen by a panel of almost 500 fungal pathologists [Dean et al., 2012] as it causes up to 30% of global rice production losses [Skamnioti and Gurr, 2009]) and *Batrachochytrium dendrobatidis* (blamed for recent decline of 500 amphibian species and complete extinction of 90 of them [Scheele et al., 2019]). The rest of the species were assigned randomly. Overall, we placed 10% of all species in test and validation sets each, while the remaining 80% formed the training set.

To model the task of predicting pathogenic potentials from NGS samples, we simulated training, validation and test reads using Mason [Holtgrewe, 2010]. First, we generated 10 million single reads, 1.25 million single reads and 1.25 million paired reads per class for the training, validation and test sets respectively, keeping the number of reads per genome proportional to each genome’s length. This mirrors the protocol used for bacteria [Bartoszewicz et al., 2020] and viruses [Bartoszewicz et al., 2021b], resulting in equal mean coverage for all genomes. A side effect of such an approach is that reads originating from longer genomes could be over-represented compared to those from shorter or incomplete genomes, possibly causing generalization problems for trained machine learning models. To tackle this issue, we also simulated a second version of the training set, where the number of reads per genome is proportional to a logarithm of a given genome’s length. We call this version of the training set the ‘logarithmic-size’ set to differentiate it from the previous (‘linear-size’) approach. Note that while the ‘logarithmic-size’ training set may help balance the resulting classifiers’ performance, in a real sample, we expect the coverage to be approximately equal for equally abundant species. Therefore, we only used the ‘linear-size’ versions of validation and test sets. In both cases, the datasets maintain the 8-1-1 proportions on both read and genome level, with a total count of 25 million reads per class.

A crucial difference compared to previous pathogenic potential prediction studies is that fungal genomes are orders of magnitude longer than bacterial or viral ones. While the procedure mentioned above produced a mean coverage of 1.82 for the bacterial dataset presented in [Bartoszewicz et al., 2020], it results in a mean coverage of only 0.15 on our data. We expected this could cause training problems, as the machine learning models would only have access to a small fraction of the overall sequence diversity of the training set. Therefore, we also simulated ‘high-coverage’ versions of both the ‘linear-size’ and ‘logarithmic-size’ training sets. In this setup, we increased the total number of training reads to 240 million, compared to 20 million for the ‘low-coverage’ versions described before. The ‘high-coverage’ versions keep the mean coverage of 1.82. As the reads were simulated randomly, we expected the ‘low-coverage’ validation and test sets to be representative, and correctly model a common case of low abundance of the target pathogen compared to other DNA sources in the sample (e.g. the host). We therefore used only ‘low-coverage’ validation and test sets. In summary, we generated four versions of the training read set and one version of the validation and test set each. The datasets contain the same number of reads for each of the classes and can be easily reused for future machine learning and benchmarking applications. Finally, we simulated a temporal test set based on the genomes added to GenBank up to 12 weeks after the main part of the database was compiled (see Supplementary Note 1). We generated 625,000 reads using the ‘linear-size’ setup and balancing the number of reads per class; the average coverage matched our main test set.

#### Joint bacterial, viral and fungal datasets

As the bacterial and viral datasets [Bartoszewicz et al., 2021a] contain only 12.5 million reads per class (with the same 8-1-1 proportion of the training, validation and test sets, stratified by bacterial species or virus), we used the low-coverage versions of our ‘positive’ read sets, matching this number. We also merged the original negative classes (commensal bacteria and non-human viruses) into a joint non-pathogen class, downsampling accordingly. Further, the original datasets contain reads of between 25 and 250bp instead of 250bp only, which makes the problem more challenging, but increases robustness on very short sequences [Bartoszewicz et al., 2021a]. To make our fungal reads compatible with this setup, we randomly shortened the reads in our validation and low-coverage training sets to 25–250bp. We only used the test sets of 250bp in order to highlight the upper performance limit of the resulting classifier.

Each of the classes contains 12.5 million reads divided in the training, validation and test subsets while keeping separate species or viruses in each of the subsets. This is intended to model a scenario of analysing a clinical sample containing a mixture of previously unseen pathogens and non-pathogens, while keeping the task feasible by resigning from differentiating between bacteriophages and commensal bacteria, as well as fungi with human and non-human hosts.

### Supplementary Note 3: Prediction algorithms and hyperparameter tuning

Following the convention introduced previously for bacteria [Deneke et al., 2017] and viruses [Bartoszewicz et al., 2021b], we focus on predicting the pathogenic potential of the analyzed species, rather than the pathogenicity itself. This distinction captures the fact that the pathogenic phenotype may or may not be manifested depending on specific host-pathogen interactions; whether this happens cannot be fully predicted outside the context of a particular host. We can only predict if a given sequence originates from a species that could cause disease under some circumstances (i.e. its pathogenic potential), not whether an invasive infection actually occurred. However, the task remains inherently difficult – a DNA sequence by itself, outside of the context of the host, can be probabilistically associated with a certain trait, but the trait itself is only realized in the biological system comprising both the host and the colonizing microbe as a whole. Therefore, we intentionally include opportunistic pathogens in the ‘positive’ class of human pathogens. These considerations mirror challenges previously described for other pathogen classes [Deneke et al., 2017, Bartoszewicz et al., 2020, 2021b,a]. Given limited information content of a short NGS read, predicting if a fungus can infect humans directly (and only) from reads is therefore an extremely challenging task.

The ResNet architecture used in this work consists of 17 convolutional layers of between 64 and 512 filters (with filter size of 7 for the first layer and 5 for the following layers) using skip connections, followed by a global average pooling layer and a single-neuron output layer with sigmoid activation. It returns an output score between 0 and 1, where a threshold of 0.5 is used by default as a boundary between ‘positive’ and ‘negative’ predictions. The network uses batch normalization and input dropout, which can be understood as randomly switching a predefined fraction of input nucleotides in training samples to *N* s. It also guarantees identical predictions for any given sequence and its reverse-complement via parameter sharing. For more details, we refer the reader to the corresponding publication [Bartoszewicz et al., 2021a].

As previous research reported that the input dropout rate is an important hyperparameter [Bartoszewicz et al., 2020, 2021b,a], we investigated two different values – the default 0.25 and 0 (no dropout). To this end, we trained the networks using the ‘linear-size’ and ‘logarithmic-size’ versions of the low-coverage training set for the default maximum of 30 epochs with an early stopping patience of 10 epochs, selecting the epoch and input dropout rate resulting in the highest accuracy on the validation set. We then retrained the networks using the selected input dropout rates and both versions of the high-coverage training set. The multi-class networks use the default input dropout rate (0.25), as required by the addition of viral and bacterial data. We trained those models for 50 epochs due to the increased difficulty of the task.

As shown in [Bartoszewicz et al., 2020], predictions for two reads in a read pair can be averaged, improving prediction accuracy; training the networks on single reads allows to use them for both single reads and read pairs. Further, averaging over predictions for all reads originating from the same organism (e.g. for classification of whole genomes or single-species samples) has been demonstrated to yield accurate species-level predictions [Deneke et al., 2017, Bartoszewicz et al., 2020]. We adapted those steps to our models as well. However, since the networks are explicitly trained for read-based classification only, the default classification threshold may be suboptimal for the genome-wise classification scenario. Therefore, we hypothesized that it should be retuned for optimal performance according to a preselected metric, e.g. balanced accuracy. To test this assumption on a fully independent dataset first, we used the ‘novel viral species’ dataset from [Bartoszewicz et al., 2021b] and their convolutional neural network (CNN) without any retraining. We selected the threshold optimizing the balanced accuracy on the appropriate validation set, rounded to two decimal places. We compared the original and retuned models and applied the procedure resulting in the better test set performance to our fungal models.

To classify novel fungal sequences based on their closest taxonomic matches, we used BLAST [Altschul et al., 1990, Camacho et al., 2009] as described for bacterial data in [Bartoszewicz et al., 2020]. The reference database was created from the training set genomes. We used discontiguous megablast with an E-value cutoff of 10 and default parameters, selecting the label of the top hit (‘pathogenic to humans’ or not) as a predicted label for each query sequence. Note that if no hits are found, no prediction can be returned for a given read. This lowers the true positive and true negative rates, defined as the ratios of correct positive or negative predictions to all reads in the positive or negative class, respectively. For read pairs, a single match is enough to assign a label to a pair, but conflicting matches result in no prediction. Similarly to [Bartoszewicz et al., 2021b], we considered two approaches for generating species-level predictions. First, we found BLAST matches of contigs from the test set, using a majority vote over all contigs belonging to the same species to assign a corresponding label. Second, we used the majority vote over predictions for all reads originating from a given species. The latter case is especially relevant for a use-case of a sequencing sample containing a single-species, and is more directly comparable to the genome-level evaluation of the ResNet.

**Table S2:**
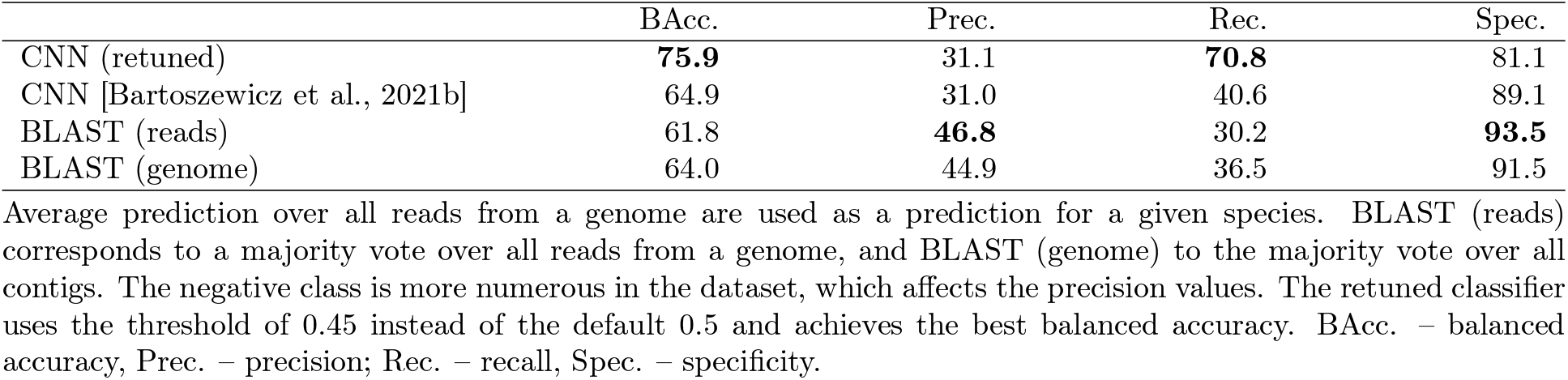
Classification threshold tuning for full available genomes on the ‘novel viral species’ dataset of DeePaC-vir [Bartoszewicz et al., 2021b].

### Supplementary Note 4: Relationship between *s*(*E*[*z*]) and *E*[*s*(*z*)]

#### Methods

Averaging the activations of the penultimate layer for a set of reads belonging to the same genome and passing the output to the final layer, we obtain the value *s*(*E*[*z*]), where *s*(*x*) = 1*/*(1 + *e*^*−x*^) is the sigmoid activation and *z* = **w**^*T*^**h** + *b*, for the penultimate activation vector **h**, weights of the last layer **w** and its bias *b*. Note that this is *not* mathematically equivalent to *E*[*s*(*z*)], or the expected output of the network for reads originating from a given genome, which was used for the genome-level predictions in previously published literature [Bartoszewicz et al., 2020, 2021b], as well as in this study. However, we suspected that although usually *s*(*E*[*z*]) and *E*[*s*(*z*)] are not identical, those terms could have similar values in practice, as they both express a notion of aggregating information contained in individual reads to yield a prediction for a full genome. If this is the case, mean activations for a set of reads originating from the same genome can be used as if they were the internal genome representations of our species-level classifiers, even though the models rely on aggregating read-level predictions. To investigate the relationship between *s*(*E*[*z*]) and *E*[*s*(*z*)], we performed additional experiments.

We speculated that average activations of the penultimate layer *E*[**h**] could be used as representations of full genomes, even though our genome-level classifiers do not explicitly use *E*[**h**], but rely on average per-read predictions *E*[*s*(*z*)]. Note that *E*[*s*(*z*)] represents averaging in in the space of predicted probabilities (‘proba’-average), while *s*(*E*[*z*]) is averaging in the logit space (‘logit’-average). To check if *s*(*E*[*z*]) ≈ *E*[*s*(*z*)], we plotted the ‘logit’-average predictions for each species against their ‘proba’-average equivalents. As any effects found could be dataset-dependent, we perform this not only on the fungal validation and test sets, but the ‘novel viruses’ dataset used for the DeePaC-vir networks (and consequently, also for our multi-class classifiers). Further, we use simulated logit values (*z*) to gain deeper insight into the relationship between both versions of read averaging. To accurately model the problem of aggregating predictions for individual species in the context of binary classifications, we first defined two classes, ‘0’ and ‘1’. We then simulated 100 ‘species’ belonging to each class. Each species *S* is described by a distribution of logit values 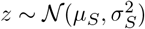, where *µ*_*S*_ and *σ*_*S*_ are the mean and standard deviation of *z* for that particular species and *𝒩* is the normal distribution. To generate *µ*_*S*_ and *σ*_*S*_ for each species, we sample from additional distributions: 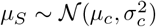 and 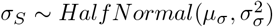, where *Half Normal* is the half-normal distribution and *c* is the class index for classes ‘0’ and ‘1’. Note that for simplicity, we sample *σ*_*S*_ from the same distribution irrespective of the species’s class. Finally we generate 1000 values of *z* per species (corresponding to 1000 reads) and plot the ‘logit’-averages *s*(*E*[*z*])] against the ‘proba’-averages *E*[*s*(*z*)]. Hence, to each species belonging to a given class, we assign its own mean and standard deviation used to generate read representations according to the species’s class. First, we perform two experiments using the following parameters:

- *µ*_0_ = −3, *µ*_1_ = 3, *σ*_0_ = *σ*_1_ = 2, *µ*_*σ*_ = 0.25, *σ*_*σ*_ = 0.25
- *µ*_0_ = −3, *µ*_1_ = 3, *σ*_0_ = *σ*_1_ = 2, *µ*_*σ*_ = 3, *σ*_*σ*_ = 1

This models two well-separated classes, and the only difference between the two settings is the distribution of within-species variances of *z*.

We then test if the relationship between ‘logit’- and ‘proba’-average changes if no separable classes are present in the dataset. To this end, we perform two additional experiments with only one ‘class’, ‘0’:

- *µ*_0_ = 0, *σ*_0_ = 3, *µ*_*σ*_ = 0.25, *σ*_*σ*_ = 0.25
- *µ*_0_ = 0, *σ*_0_ = 3, *µ*_*σ*_ = 3, *σ*_*σ*_ = 1

Finally, we investigate if any effects found could be conditional on the presence of a species-level signal, i.e. on differences in the distribution of *z* for each species *S*. We generate 100,000 values of 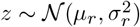, where *µ*_*r*_ = 0 and *σ*_*r*_ = 20, and assign the ‘reads’ to 100 simulated ‘species’ arbitrarily (first 1000 reads to the first species, second 1000 reads to the second species, etc.). We also repeat the experiment simulating 5000 ‘reads’ per ‘species’ (corresponding to a 5-fold increase in coverage).

#### Results

As shown in Figure S2a and Figure S2b, the values of *s*(*E*[*z*]) and *E*[*s*(*z*)] are indeed similar for both the validation and test set. They are highly correlated (Spearman’s *ρ >* 0.99, *p <* 10^*−*15^) and linearily dependent. This justifies using *E*[**h**] as our genome representations. However, for the ‘novel viruses’ dataset, the relationship actually resembles a sigmoid function (Figure S2c). To investigate the relationship between the two versions of read averaging, we performed further experiments using simulated logit values *z*. In the binary classification case, if *σ*_*S*_ is low, the relationship between ‘logit’- and ‘proba’-averages is linear (Figure S3a), and if *σ*_*S*_ is high, the relationship is sigmoid (Figure S3b). We suggest an intuitive explanation of this effect. High within-species variance leads to many large absolute values of *z* within a given species. Note that extreme outliers disproportionately influence the mean, or *E*[*z*]. The sigmoid function applied *before* averaging ‘squashes’ those extreme values, diminishing their influence. This explains why *s*(*E*[*z*]) values tend to be more extreme than *E*[*s*(*z*)]). The effect is stable even if no separable classes are present in the dataset. Strikingly, we can easily reproduce the linear (Figure S3c) and sigmoid (Figure S3d) behaviour just by manipulating the within-species variance.

What is more, the effect requires species-level signal to be present, i.e. each species must have its own mean and standard deviation assigned. If the distributions of *z* are the same for each species, the differences in *E*[*s*(*z*)] are expected to be minimal. However, if the variance of *z* is extremely high, sampling effects can cause large dispersion of *s*(*E*[*z*]). As shown in Figure S3e, ‘logit’-averaging indeed causes spurious, high-confidence predictions in this setting (since *µ*_*r*_ = 0, correct predictions should be close to 0.5). Those are most probably artifacts caused by the extreme values of some *z* outliers. ‘Proba’-averaging is more robust in this case and correctly yields low confidence predictions close to 0.5. Note that this can be also understood as the ‘logit’-averaging amplifying noise generated by sampling effects. Therefore, the problem can be at least partially mitigated with higher read numbers per genome – if we increase the number of ‘reads’ per ‘species’, we see the expected decrease in effect strength (Figure S3f). In general, higher sensitivity of ‘logit’-averaging could also be useful in some applications, depending on the levels of different sources of variance in the dataset.

Note that even in the settings with high within-species variance, where the relationship between *s*(*E*[*z*]) and *E*[*s*(*z*)] becomes sigmoid, its approximate monotonicity is maintained (Spearman’s *ρ >* 0.98, *p <* 10^*−*15^ for both Figure S3b and Figure S3d). This suggests that even in this case, *E*[**h**] could potentially be used as genome representations. However, one must exercise caution – the more steep the sigmoid relationship, the more distorted the distances between such representations would become. In our fungal dataset, the within-species variance is seemingly low enough to guarantee a linear relationship, so this problem does not affect it.

**Figure S2:**
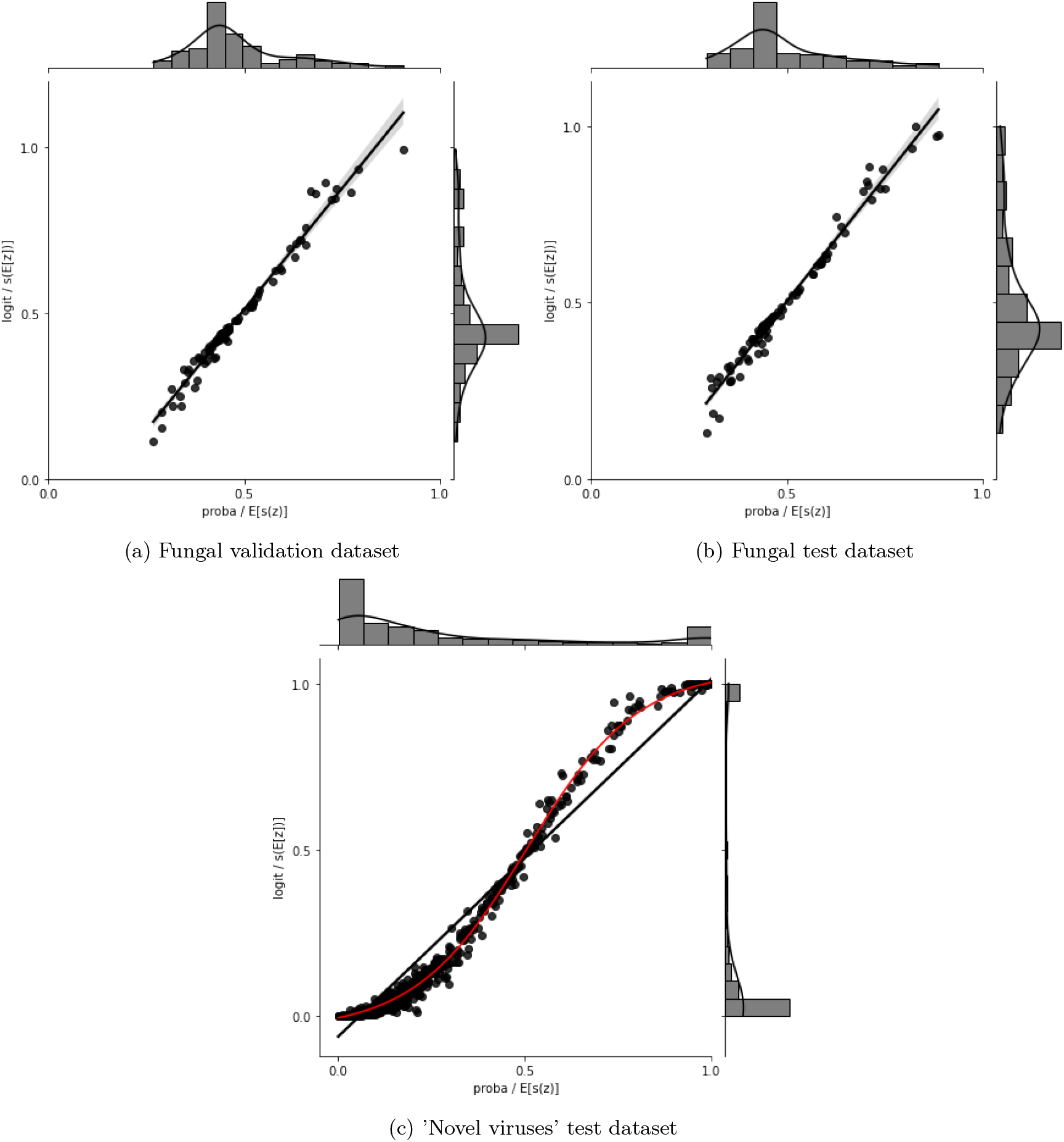
Comparison of relationships between ‘logit’-average (*s*(*E*[*z*]) and ‘proba’-average (*E*[*s*(*z*)]) predictions. Real data.

**Figure S3:**
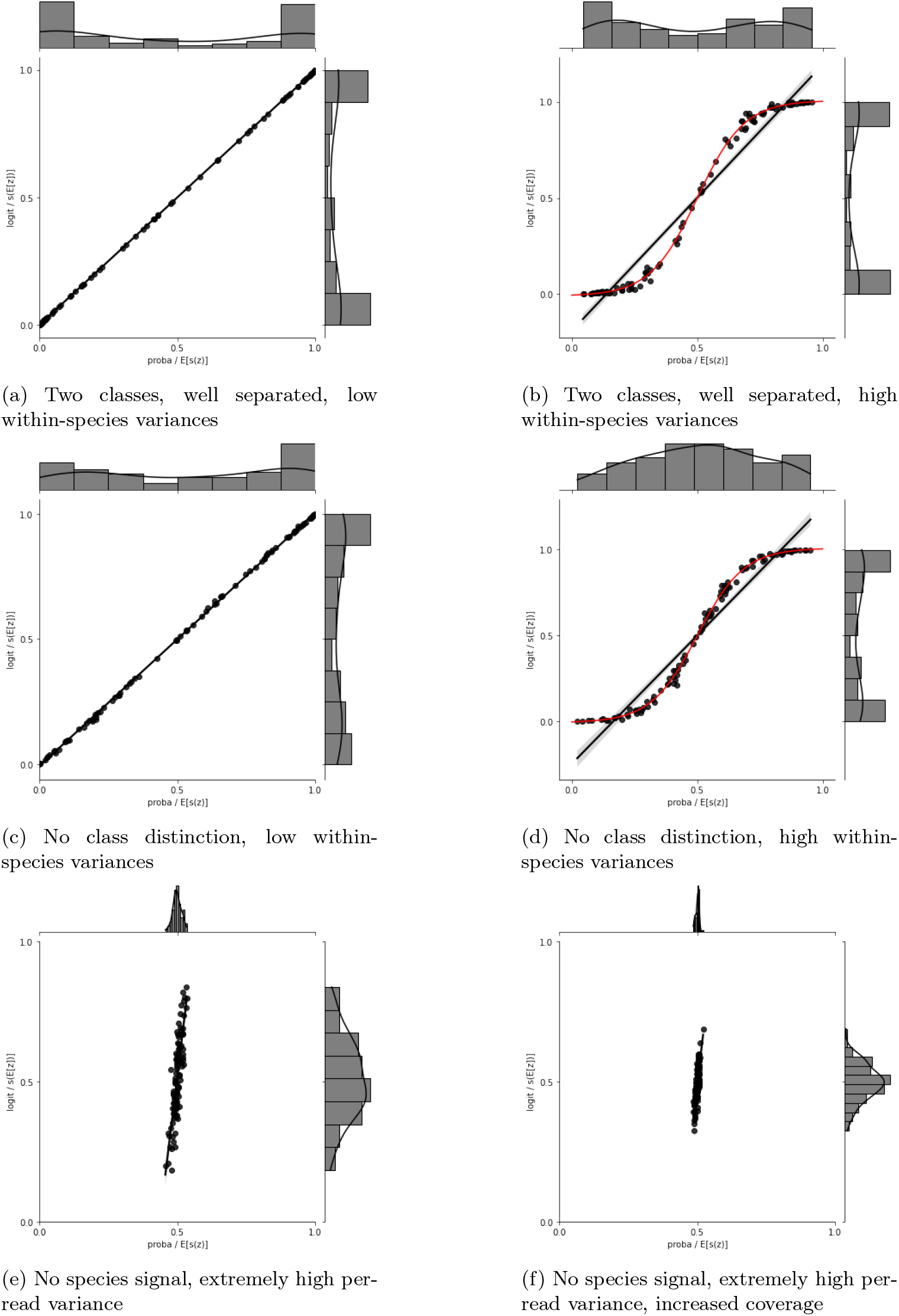
Comparison of relationships between ‘logit’-average (*s*(*E*[*z*]) and ‘proba’-average (*E*[*s*(*z*)]) predictions. Simulated *z* values.

### Supplementary Note 5: Fungal pathogenic potential prediction

The networks trained without input dropout achieved higher validation accuracy compared to those trained with dropout, and the ‘logarithmic-size’ training set resulted in higher accuracy than the ‘linear-size’ set in all cases. As expected, the ‘high-coverage’ training set also improved performance, although the training time of around 8h45’ per epoch on four Tesla V100 GPUs was much higher than around 45 minutes per epoch for the ‘low-coverage’ variant on the same hardware.

The difference in performance for the first and second mate of the read pairs is negligible; we present the mean values. Interestingly, while BLAST’s accuracy on the test set is higher, its accuracy on the validation set is lower by a similar margin (61.6%, compared to 63.3% for the ResNet). While this comparison should not be overinterpreted, as the ResNet epoch maximizing validation accuracy was chosen, those results suggest that the performance differences between the two approaches are small and could be dependent on a particular composition of species in the test set. Although this could be explicitly tested using a nested cross-validation scheme, such a setup would be computationally prohibitive.

We do not observe much overfitting (the single read training accuracy for the selected epoch is 68.9%), which suggests that the unsatisfactory performance is a form of underfitting. More expressive neural architectures could possibly help alleviate this problem, but it is also possible that prediction accuracy known for other pathogen classes is simply impossible for fungi, for example due to the higher complexity of eukaryotic genomes with abundant regions of non-coding DNA. However, we speculate that even those predictions could be useful in some circumstances. High precision of BLAST proves that its positive predictions are trustworthy. On the other hand, the ResNet offers only slightly worse accuracy, but for all reads in the sample (BLAST finds no matches for over 20% of reads even if read pairs are used). What is more, the ResNet can process 1.25 million reads in just 3 minutes on a single GPU, while BLAST needs over three hours and 256 threads on a high-performance computing node for the same task. The ResNet is slower when used on CPUs, but still outperforms BLAST more than sevenfold. This is very important in practice, as screening of large NGS datasets must be performed quickly to remain feasible. For this reason, BLAST is usually too slow for NGS analysis. Note that in this work, it represents the upper bound on accuracy of homology-based approaches; faster alternatives like NGS mappers or k-mer apporaches have been shown to underperform in the pathogenic potential tasks for other pathogen groups [Deneke et al., 2017, Bartoszewicz et al., 2021b].

Interestingly, BLAST’s precision is actually a bit lower for full genomes than for reads, while the ResNet becomes more precise when more information (more reads) is considered. Contrary to what was reported for viruses, using raw, unassembled reads improves BLAST’s performance compared to relying on assembled contigs. A probable reason is that when a majority vote over all contigs is taken, all contigs have the same voting weight, which means that the ratio of voting weight to sequence length can be much higher for short contigs of potentially lower quality than for the very long ones. In contrast, when reads are used, all regions of the genome weight the same. Using contigs also markedly increases the processing time. This slowdown is probably best explained by the computational difficulty of the local alignment of long contigs compared to a low-coverage set of representative reads.

Both BLAST and the ResNet correctly predict *C. auris* and *A. fumigatus* to be pathogens, even though they were kept in the held-out test dataset unused during training. *P. oryzae* is also recognized as a a member of the negative class. The ResNet (but not BLAST) incorrectly classifies *B. dendrobatidis*, a deadly amphibian pathogen, as able to infect humans. This prediction could be in principle indicative of a hidden zoonotic potential that could be realized under some yet unreported circumstances (e.g. in susceptible individuals or tissues). However, without further evidence, we assume this is simply an error of the model – perfect generalization is rarely possible, even though as noted in Table S3, the prediction performance of the ResNet on full genomes is still high. It is possible that the erroneous prediction for *B. dendrobatidis* is caused by underrepresentation of related species in the training set, as the core database contains only two other species of the same order, *Chytridiomycota*, hindering the classifier from correctly modelling their pathogenic potential.

**Table S3:**
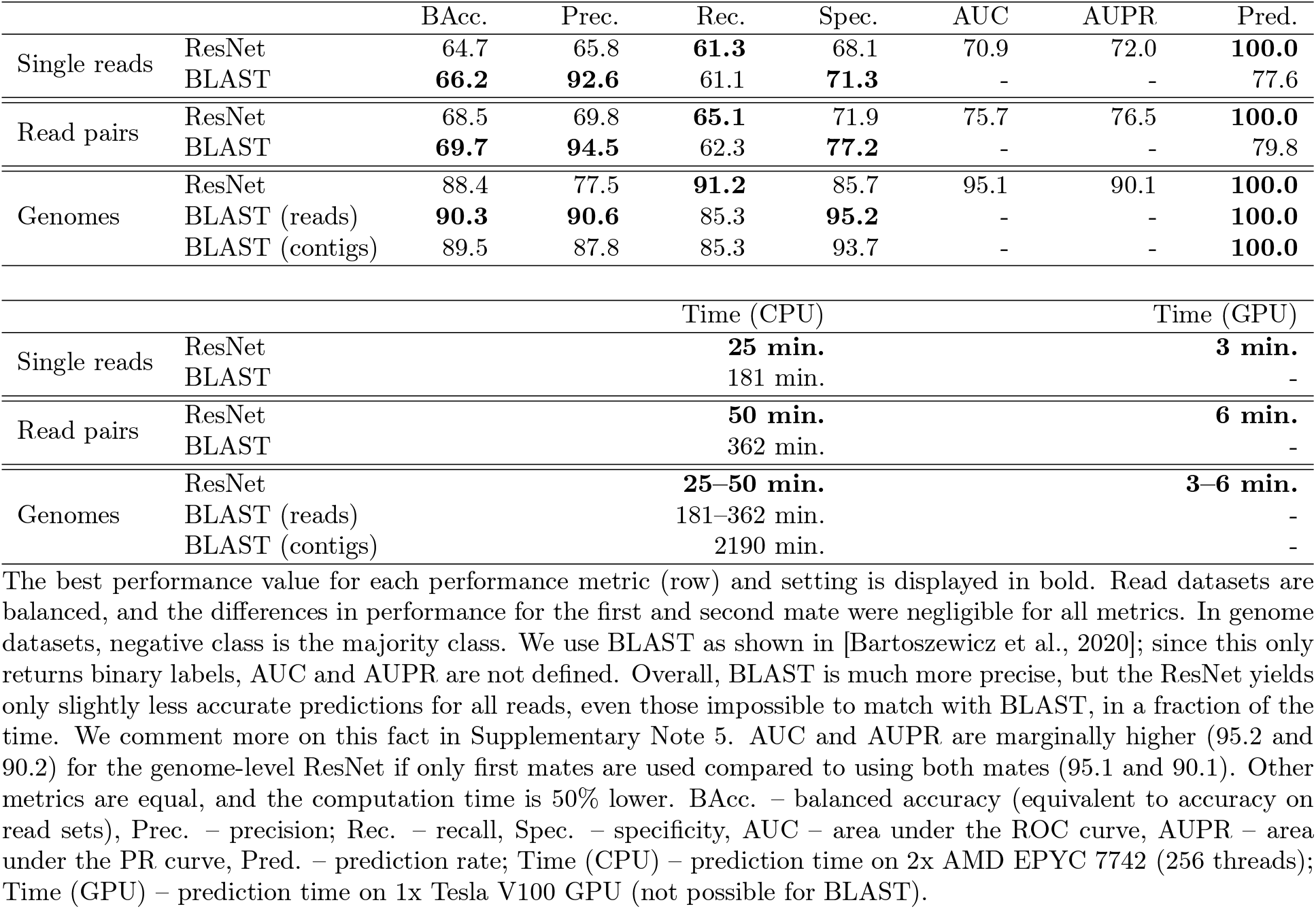
Classification performance comparison between BLAST and the ResNet model on single reads, read pairs and genomes from the test dataset.

**Table S4:**
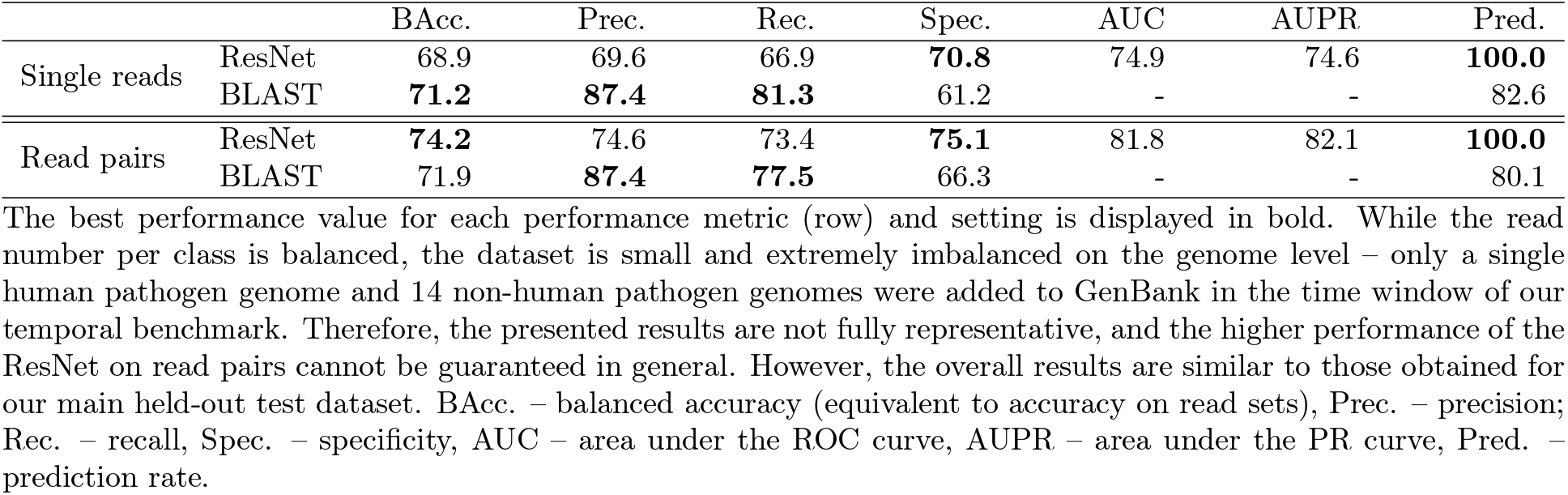
Classification performance comparison between BLAST and the ResNet model on single reads and read pairs from the temporal dataset.

**Table S5:**
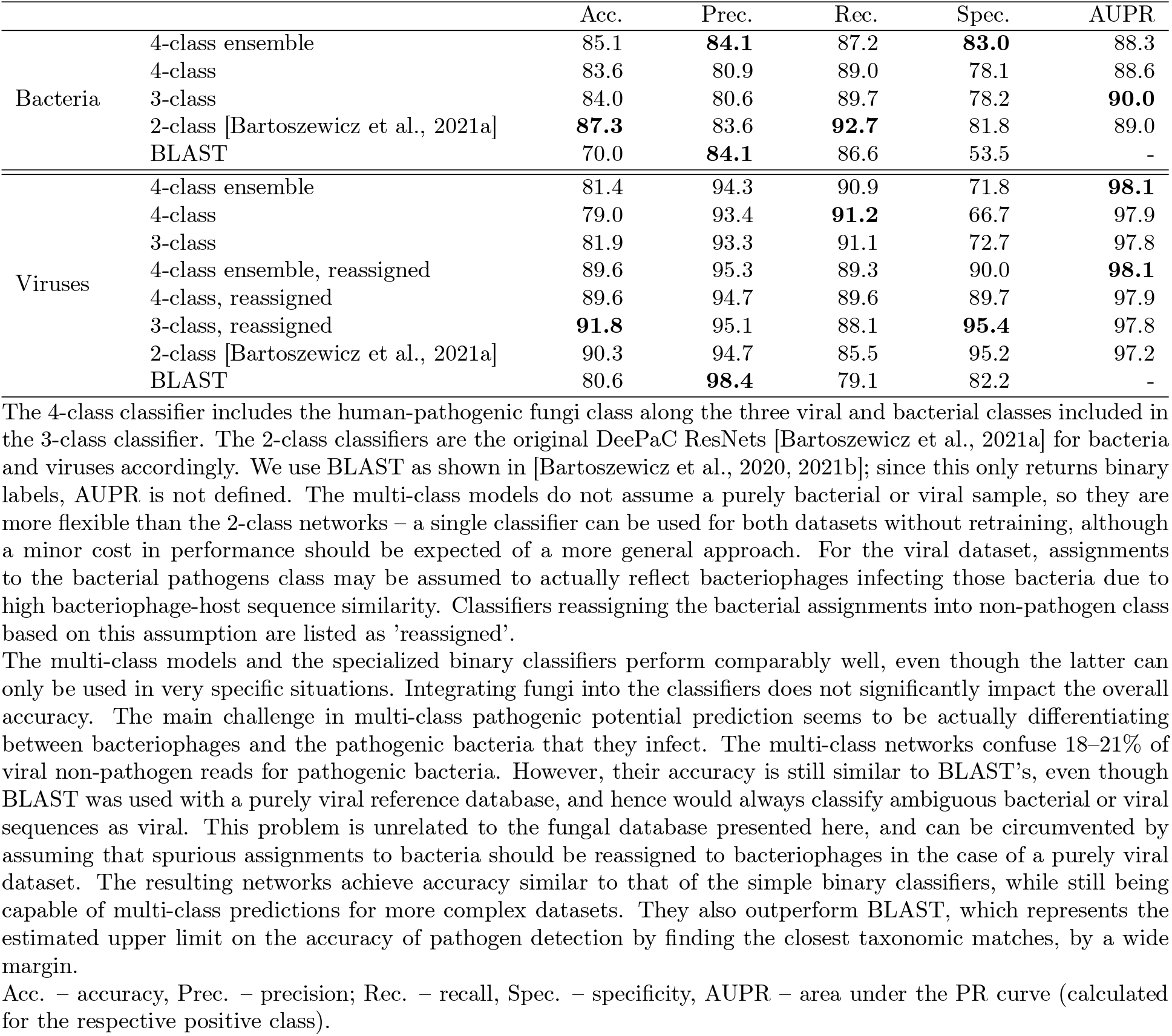
Effects of integrating multiple classes on the classification performance on non-fungal datasets, read pairs.

**Table S6:**
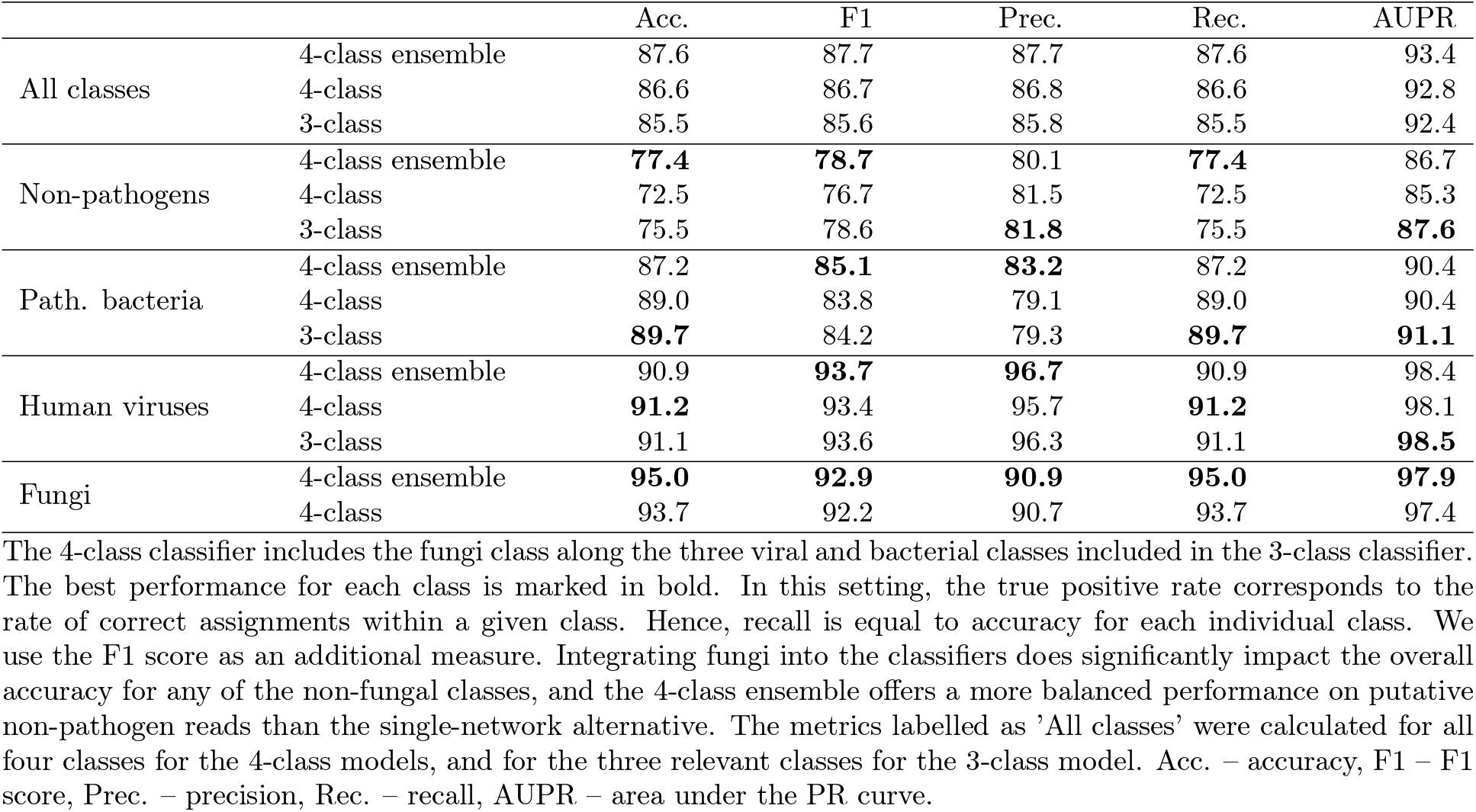
Effects of integrating multiple classes on the classification performance on the multi-class dataset, read pairs.

### Supplementary Note 6: Genome embeddings

The last convolutional layer of our fungal ResNet has 512 filters, but we removed the filters with activations equal to zero for all samples in the dataset. We then used UMAP [McInnes et al., 2020] to embed the resulting representations in a 2-dimensional space for visualization. We used the Euclidean distance metric with a minimum possible distance of 0.1 and the neighbourhood size of 15. The inputs had been randomly shuffled beforehand to avoid artifacts that can appear if an embedding is learned based on representations ordered by class.

The number of units at most levels of taxonomy poses a challenge for intuitive visualization of the dataset structure. Nevertheless, some correspondence between cluster membership, label and the taxonomic rank of a class is clearly visible (Figure S7; note that the ‘class’ as a taxonomic term is distinct from the concept of a positive or negative class in machine learning). This extends also to the lower and higher taxonomic ranks of order and phylum, respectively (Figure S8-Figure S9).

Some orders of one class may belong to different clusters. For example, the orders *Chaetothyriales, Eurotiales* (including the genus *Aspergillus*) and *Onygenales* all belong to the class *Eurotiomycetes* (Figure S7, *in red), but are members of four different clusters, with Eurotiales* split into two (Figure S8). Even more strikingly, the leftmost cluster contains at least one member of all six phyla, with two most noticeable groups corresponding to the orders *Saccharomycetales* and *Mucorales* in phyla *Ascomycota* and *Mucormycota*, respectively (Figure S7-Figure S9). 89% of the members of both those orders are at least opportunistic human pathogens; this includes *Candida* species in *Saccharomycetales* and causative agents of mucormycosis, known also as the ‘black fungus’, in *Mucorales*. Therefore, we hypothesize that the grouping represents a cluster of ‘conserved’ potential pathogens, relatively easy to classify with an acceptable accuracy. A common signal at the functional level or technical artifacts could be alternative explanations; they are however less likely due to large phylogenetic distance on one side, and a consistent grouping of almost all species from those well-represented orders on the other.

**Figure S4:**
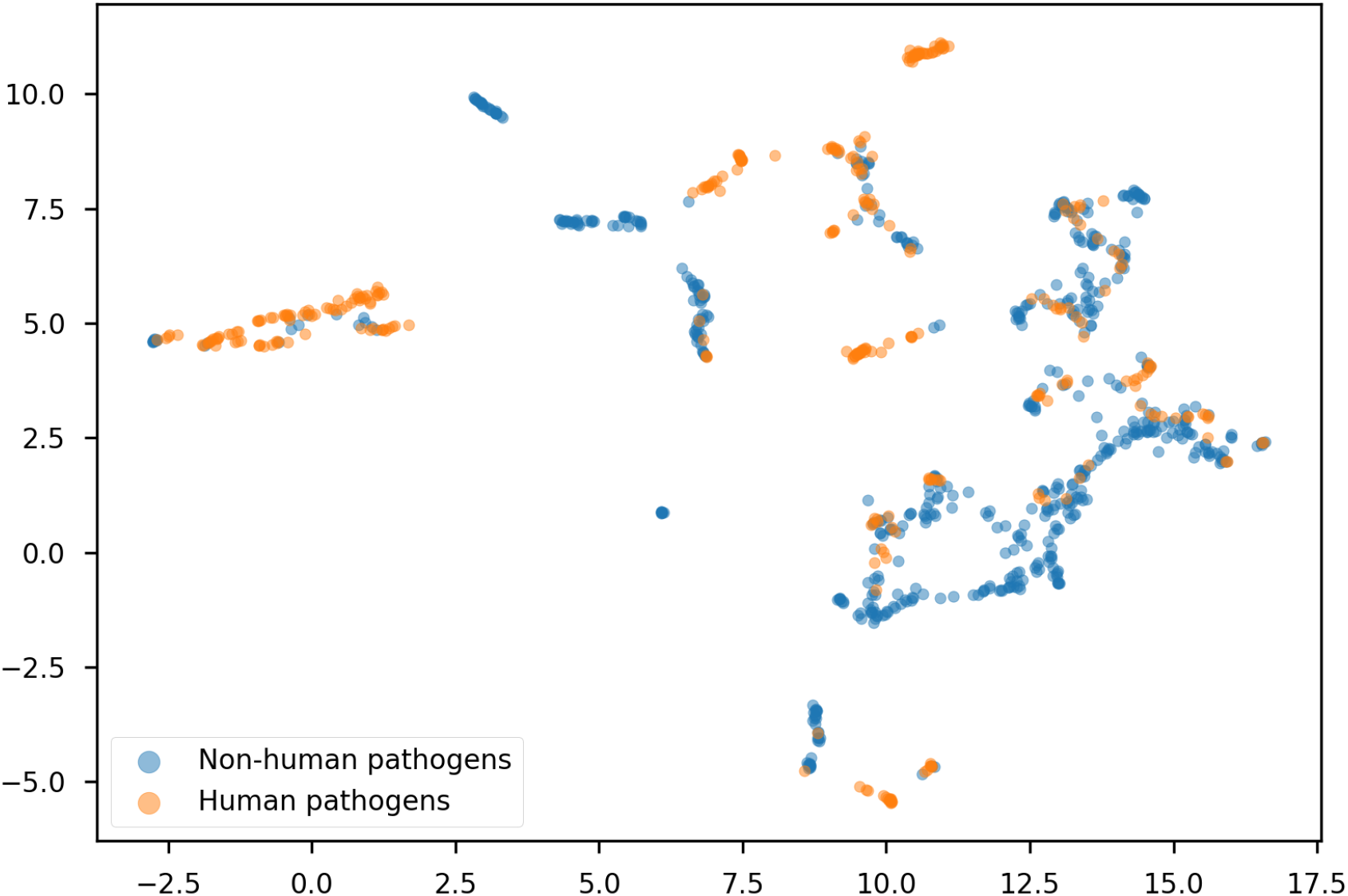
UMAP embeddings of the learned genome representations for the core database; true labels.

**Figure S5:**
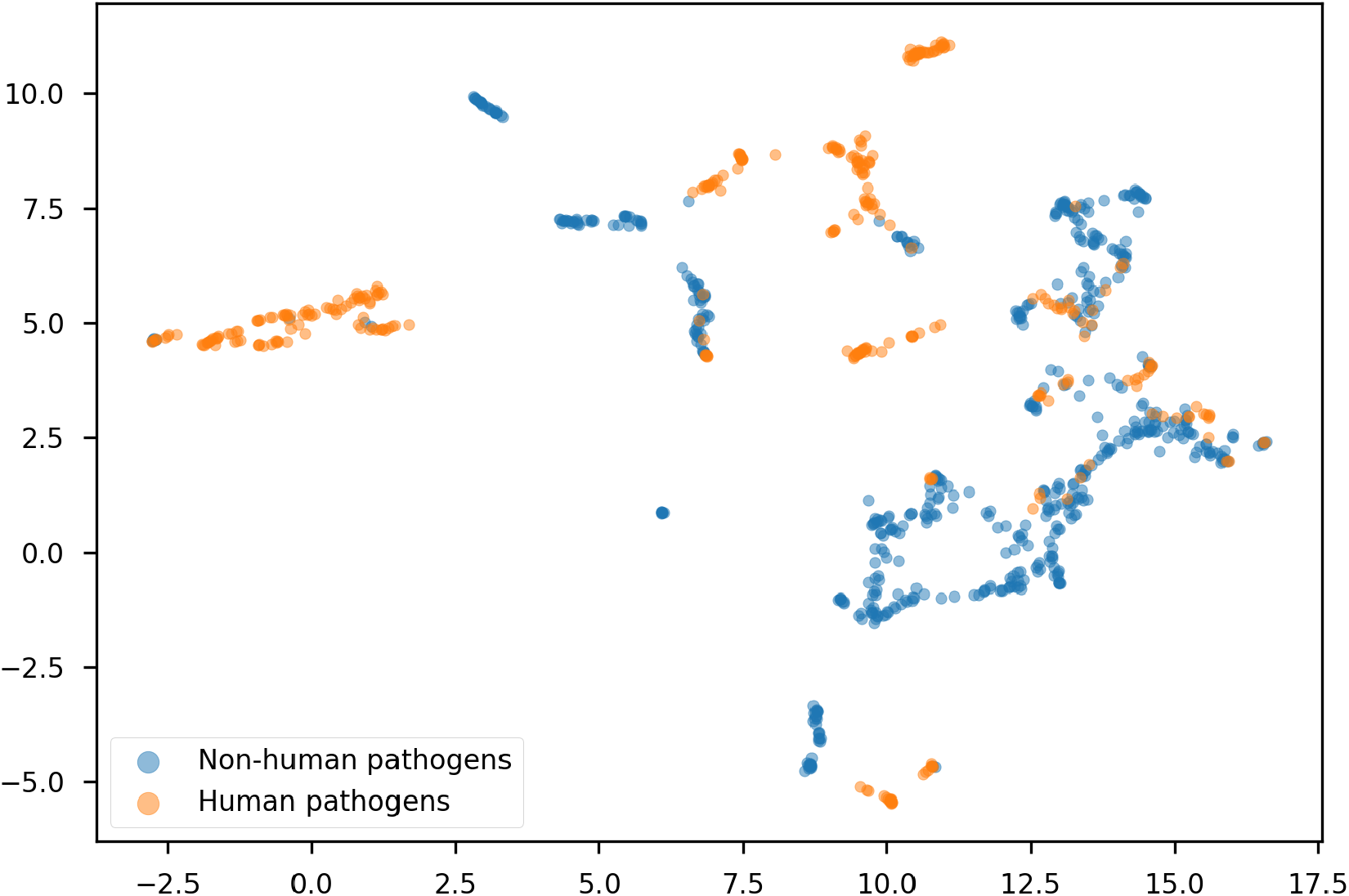
UMAP embeddings of the learned genome representations for the core database; labels predicted by the ResNet (retuned threshold). Most ‘positive’ members of otherwise ‘negative’ clusters are correctly detected.

**Figure S6:**
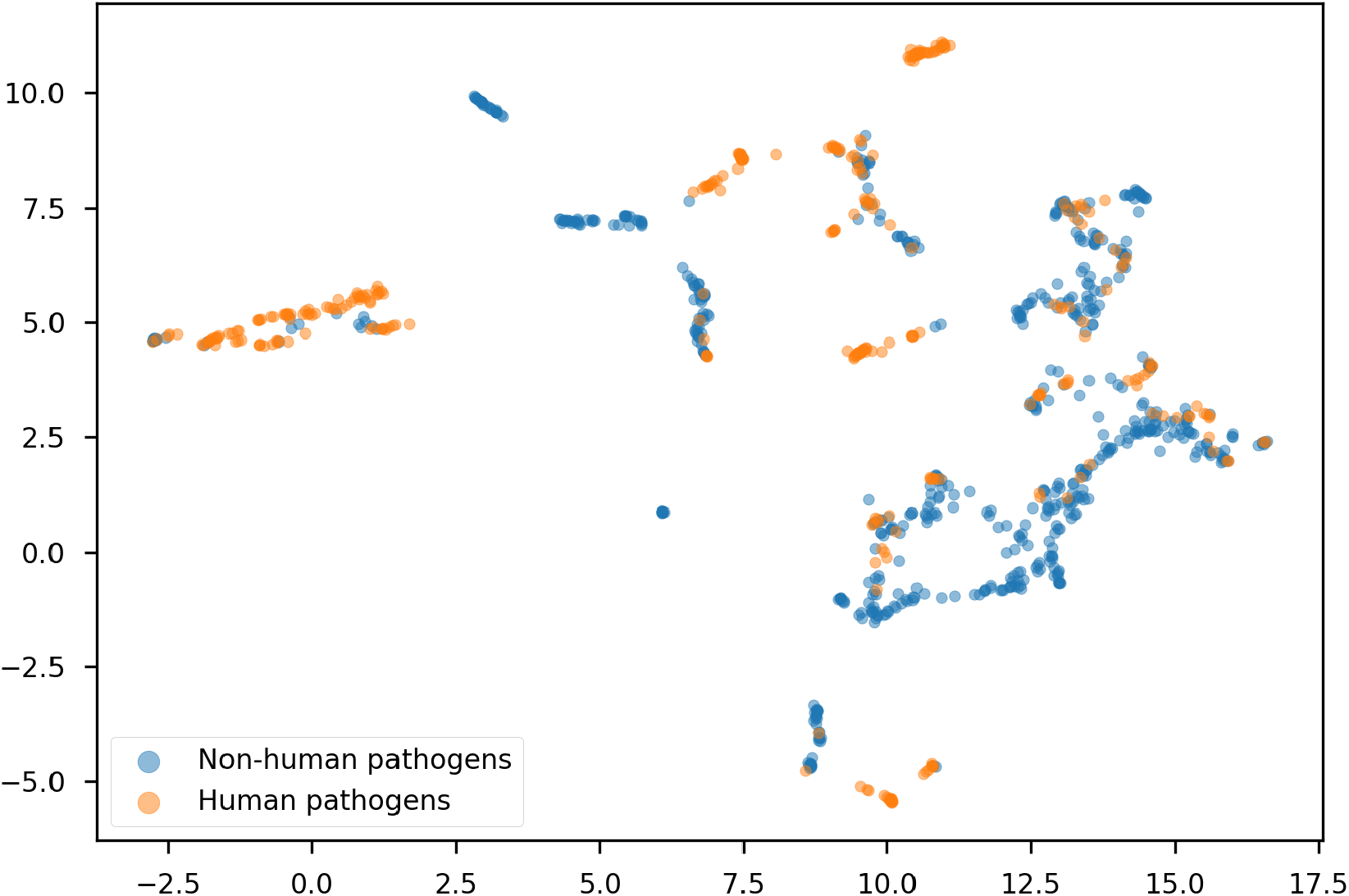
UMAP embeddings of the learned genome representations for the core database; labels predicted by BLAST. Training species are present in the reference database, so predicting labels for them is easy (99.9% accuracy).

**Figure S7:**
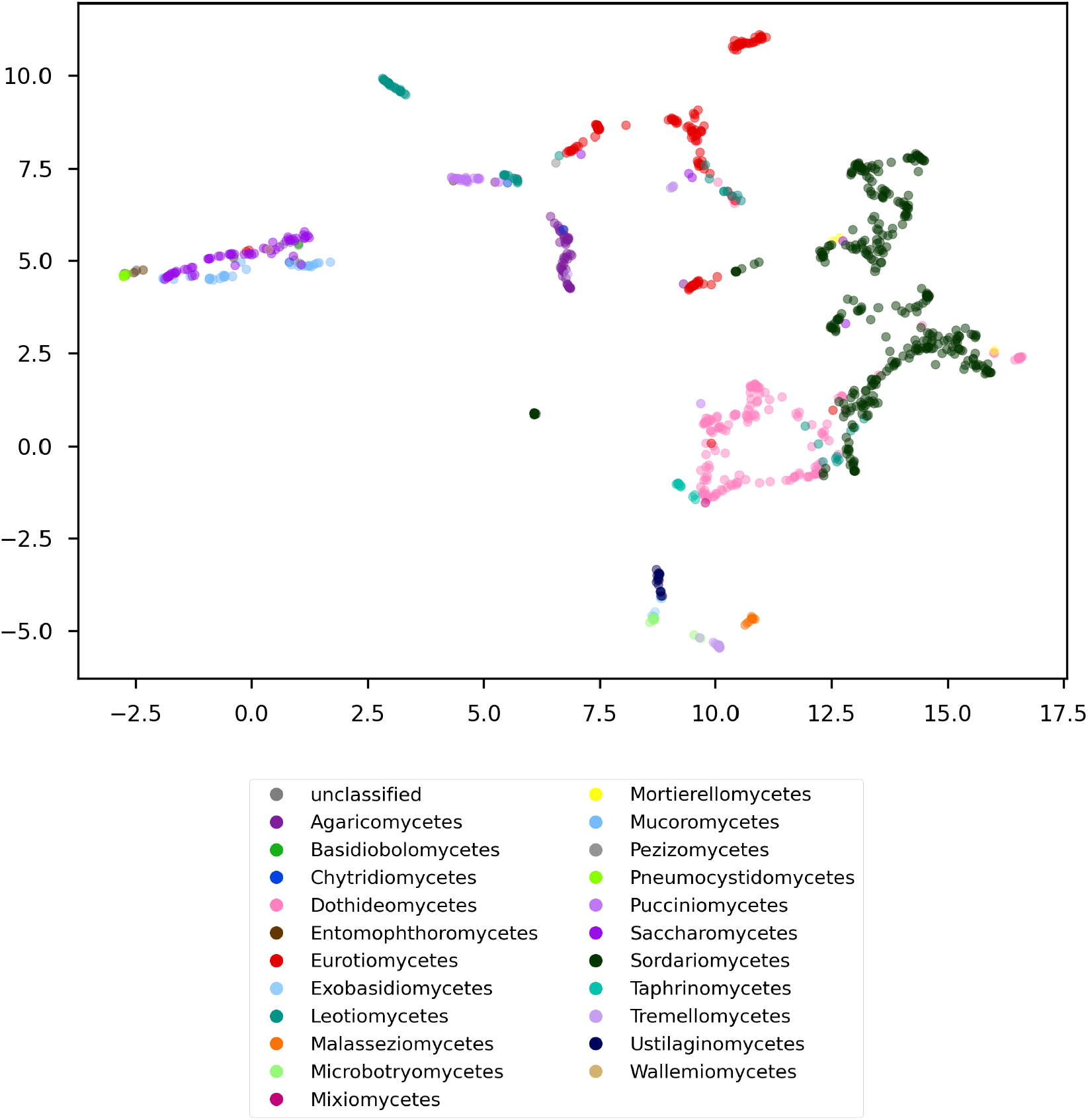
UMAP embeddings of the learned genome representations for the core database; taxonomic rank: class. In general, related species are close to each other in this space, but some taxonomic units are distributed among more than one cluster (e.g. *Eurotiomycetes*, in red), and some clusters contain members of distant taxa (e.g. the leftmost cluster).

**Figure S8:**
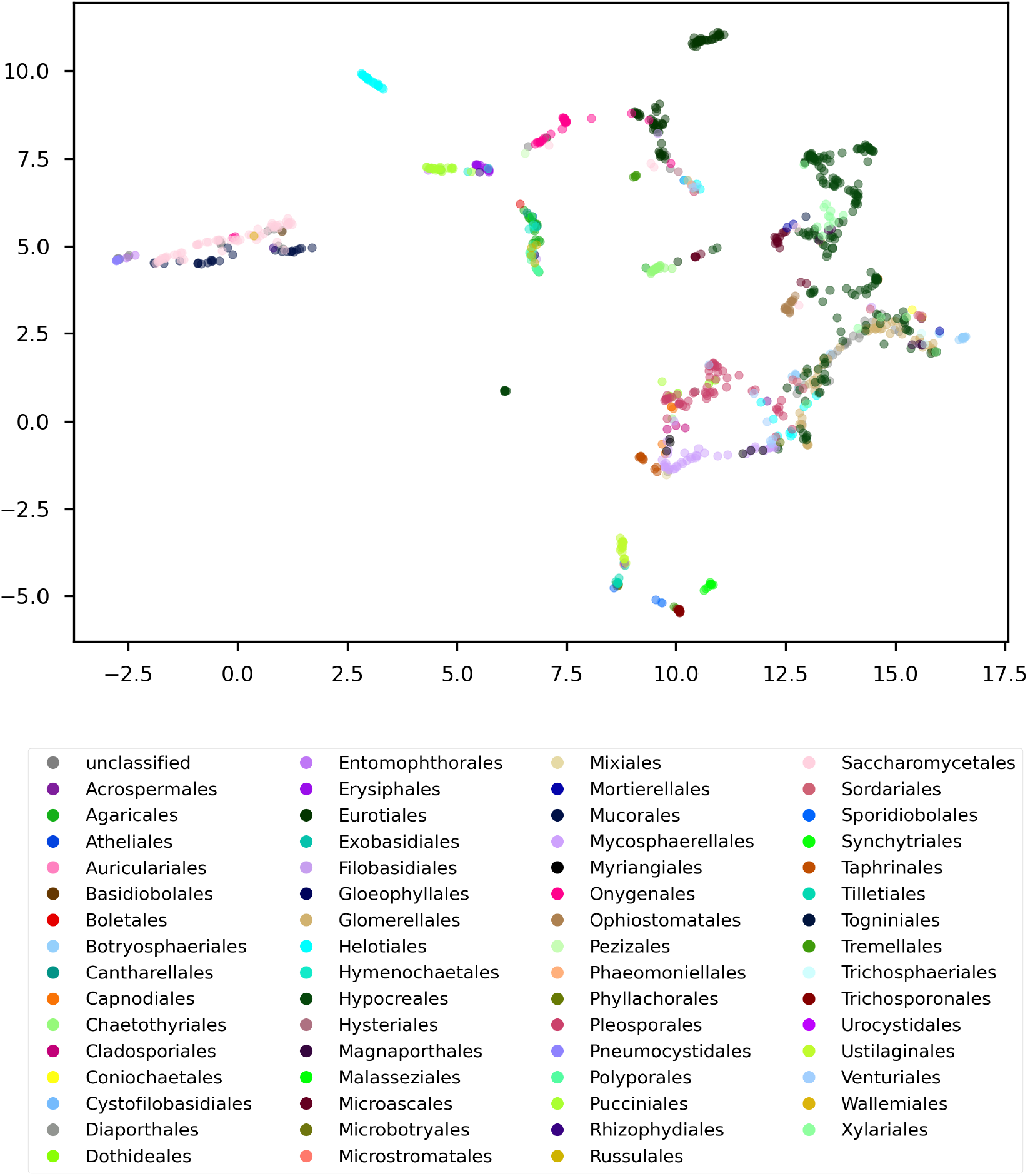
UMAP embeddings of the learned genome representations for the core database; taxonomic rank: order.

**Figure S9:**
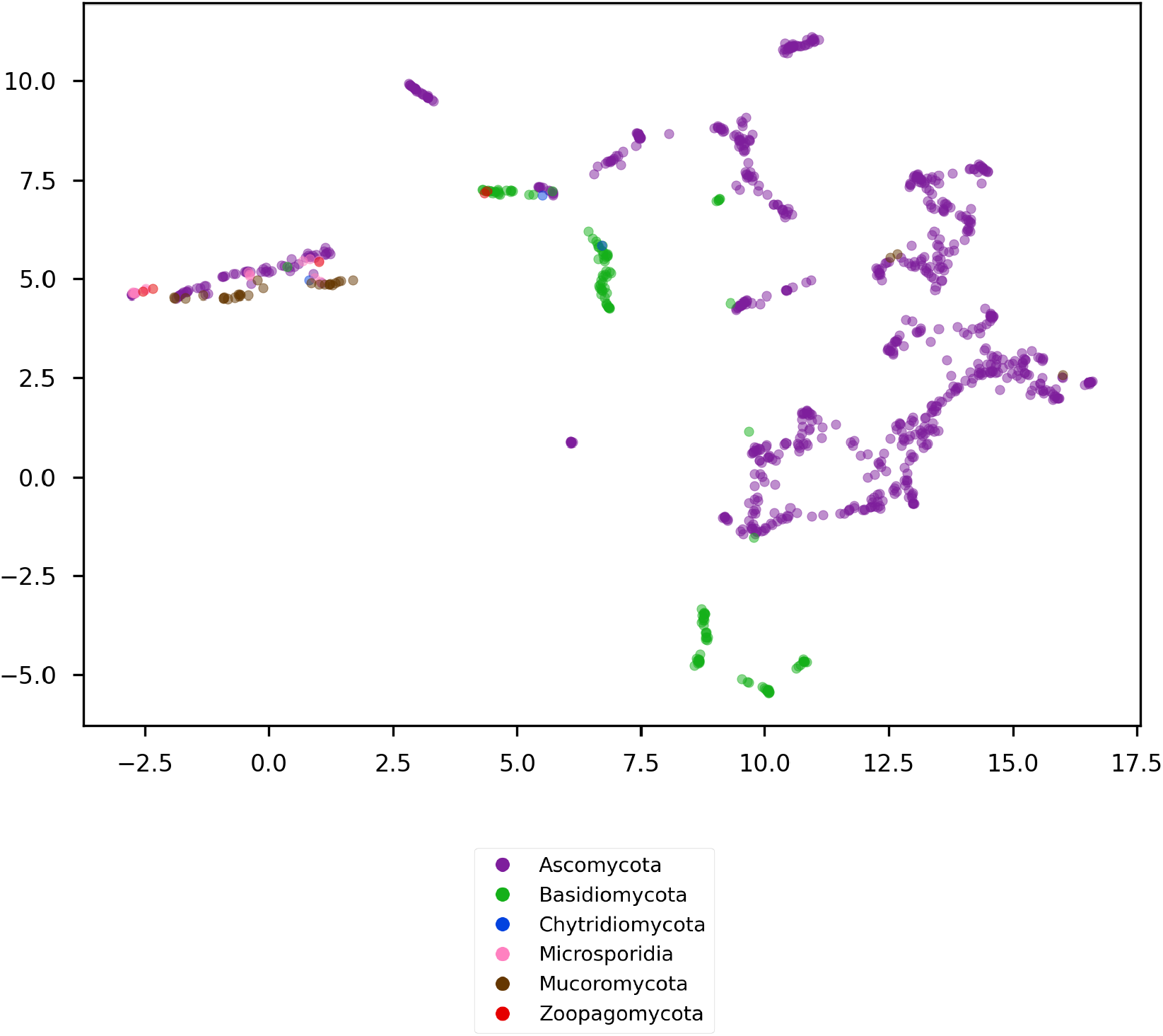
UMAP embeddings of the learned genome representations for the core database; taxonomic rank: phylum.

**Figure S10:**
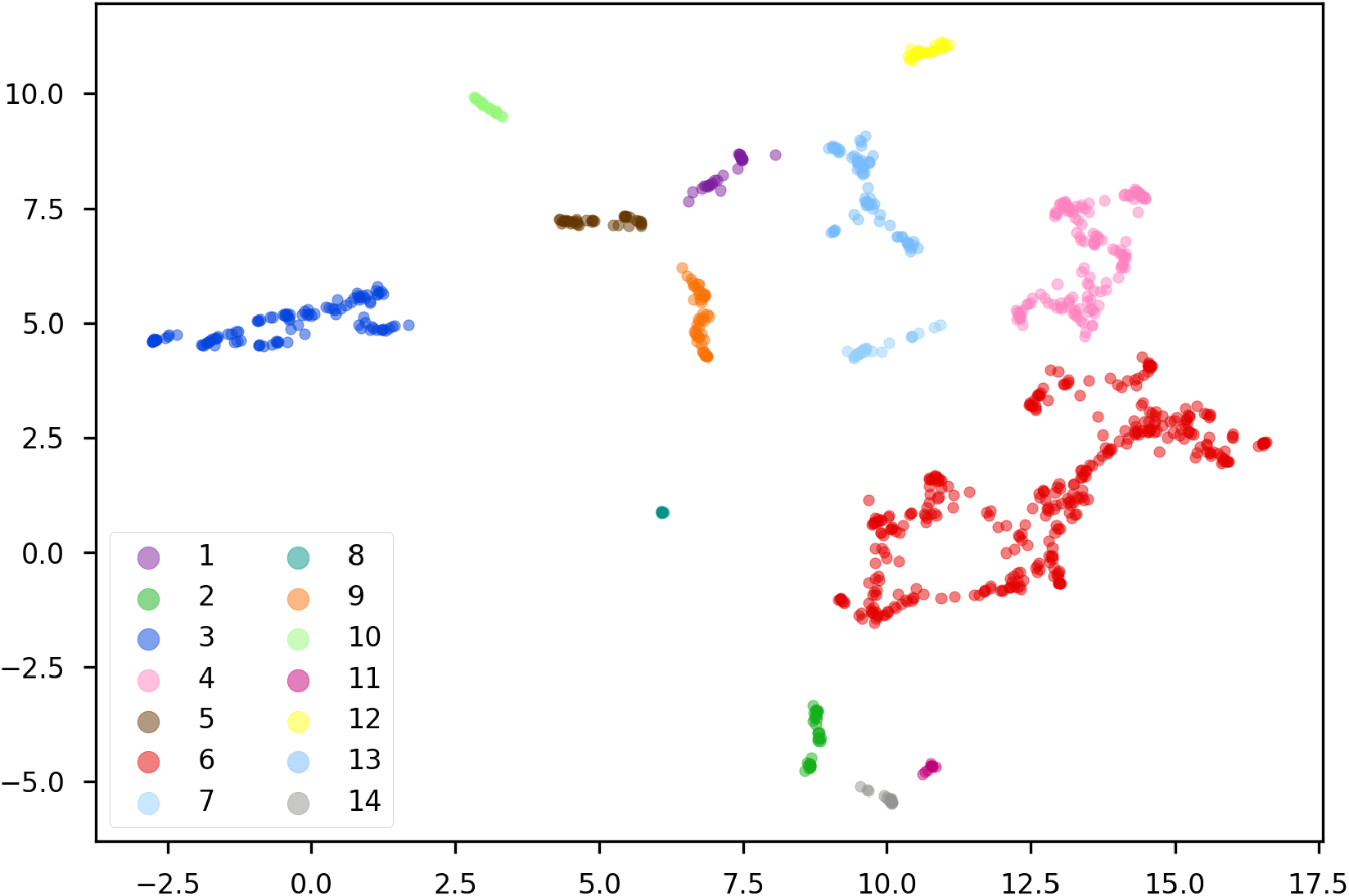
UMAP embeddings of the learned genome representations for the core database. Assignment to clusters detected with single-linkage agglomerative clustering. Clusters numbered arbitrarily.

**Table S7:**
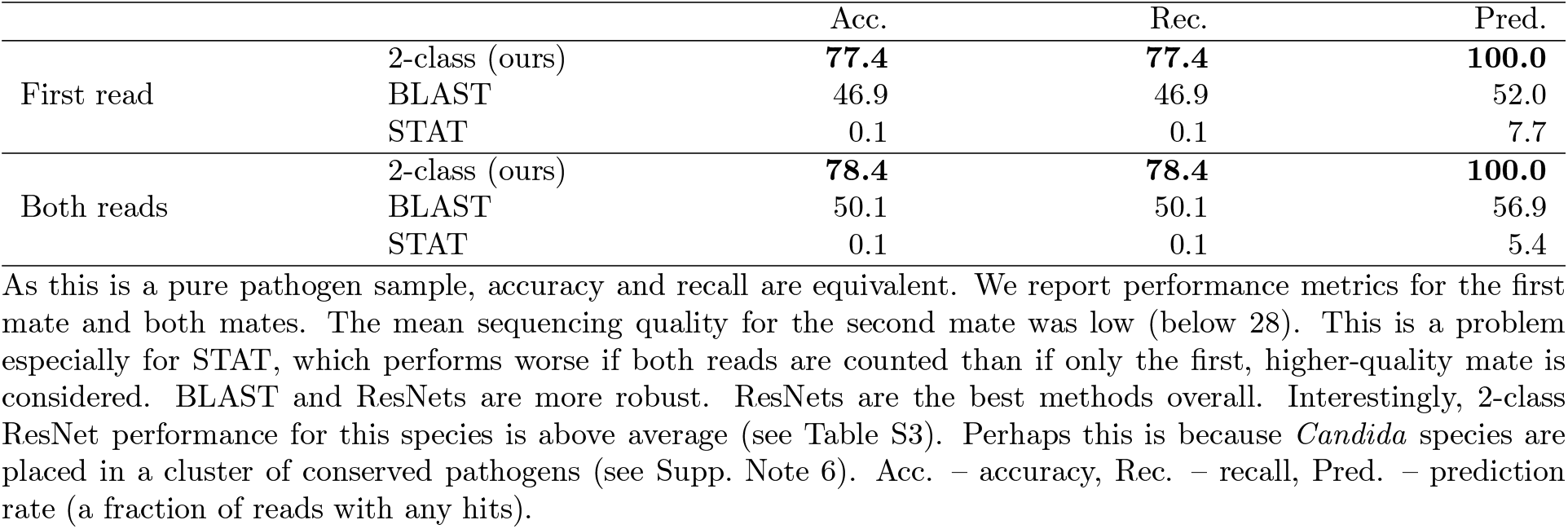
*C. auris* sequencing run, SRA accession SRR17577041. Binary classifiers (host group prediction).

### Supplementary Note 7: Performance on real data

Obtaining ground truth for genuine metagenomic samples is very challenging without spiking them with known organisms or corresponding reads. As our test sets consist of reads simulated from a mixture of different species (and viruses), they effectively function as mock metagenomic samples. Simulated reads are commonly used to evaluate computational approaches for read classification (e.g. [Wood et al., 2019, Katz et al., 2021]. To show that the methods discussed here work also on real data, we focus on a single-species sequencing run, where ground truth is available. We selected a *C. auris* sequencing run (SRA accession SRR17577041) and analyzed it in two scenarios: a) assuming it is a novel fungal species, predict whether it could infect humans (binary classification, as in Table S3-Table S4) b) assuming it is an unknown agent originating from a patient, predict its correct pathogen group (multi-class setting, as in Table 1 and Table S6). In addition to the corresponding ResNets and BLAST databases, we also used STAT [Katz et al., 2021], a k-mer tool used by the NCBI to generate taxonomy reports for all SRA archives.

To evaluate STAT, we downloaded its reference database published along the paper and run a *C. auris* strain exclusion study as described by the authors. In the host prediction setting Table S7, we counted all STAT species-level hits to any pathogenic fungal species as positives, but in the multi-class pathogen detection task (Table 2), we assumed that all reads classified as fungi can be counted as successful detection. As STAT has no way of jointly classifying read pairs, we treated both mates as independent reads also when comparing to BLAST and ResNet predictions for read pairs. Note that in contrast to our BLAST evalutation, STAT reference database contained 248,426 TaxIds (see Katz et al. [2021]), and only *C. auris* was effectively removed from it for the strain exclusion study.

