## Supplementary Information for "Detecting DNA of novel fungal pathogens using ResNets and a curated fungi-hosts data collection"

[1]Hasso Plattner Institute for Digital Engineering, Digital Engineering Faculty, University of Postdam, 14482 Potsdam, Germany; [2]Department of Mathematics and Computer Science, Free University of Berlin, 14195 Berlin, Germany. *Contributed equally.

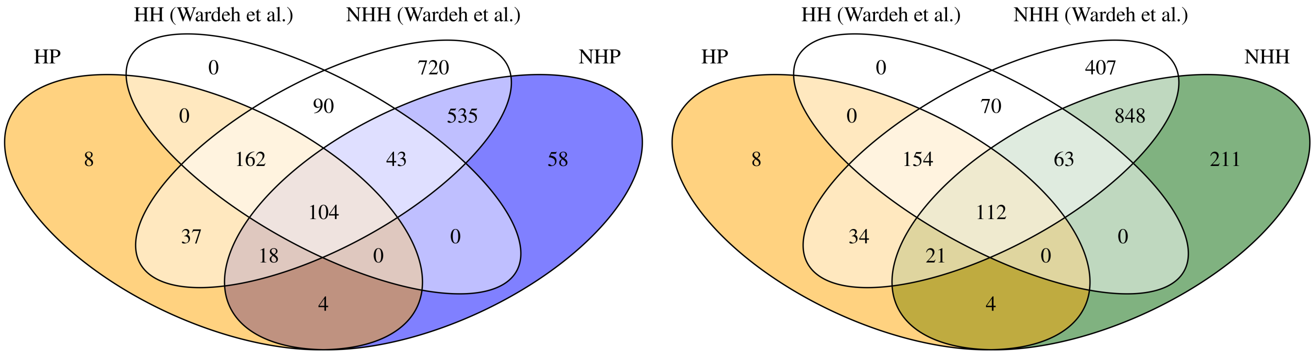


(a) Core database

(b) Core database + genomes of plant-associated species

Figure S1: A Venn diagram of the manually confirmed labels and putative labels from EID2 [Wardeh et al., 2015] for all labelled genomes (except synonyms). HP – human pathogens (manually curated); NHP – non-human pathogens (manually curated); NHH – species with a non-human host (manually curated); HH (Wardeh et al.) – species with a putative human host (EID2); NHH (Wardeh et al.) – species with a putative non-human host (EID2).

Table S1: Key information on each species included in the database.

| | |
|---|---|
| **Name** | Species name, GenBank organism name and infraspecific name for the representative genome. |
| **NCBI TaxID** | Taxonomic identifiers of the species and the representative genome in the NCBI Taxonomy database. |
| **Assembly accession & name** | GenBank assembly accession number and assembly name for the representative genome. |
| **Pathogenicity & host group** | Manually curated information regarding pathogenicity and the relevant host groups. |
| **Putative host group** | Automatically extracted information suggesting a host group. Includes non-pathogenic associations. |
| **Label sources** | References to resources used to label the species for all host groups and evidence levels. |
| **Source name and TaxID** | Species name and TaxID as mentioned in the first resource used to label it. |
| **Assembly level** | Assembly level of the representative genome. |
| **Sequence release date** | Sequence release date as reported in GenBank. |
| **FTP path** | GenBank FTP path to the representative genome. |
| **Label date** | Date of adding the species to the database or updating its labels. |

We use genomes labelled as representative or reference in GenBank.

### Supplementary Note 2: Training, validation and test read set simulation

Table S2: Classification threshold tuning for full available genomes on the 'novel viral species' dataset of DeePaC-vir [Bartoszewicz et al., 2021b].

| | BAcc. | Prec. | Rec. | Spec. |
|---|---|---|---|---|
| CNN (retuned) | **75.9** | 31.1 | **70.8** | 81.1 |
| CNN [Bartoszewicz et al., 2021b] | 64.9 | 31.0 | 40.6 | 89.1 |
| BLAST (reads) | 61.8 | **46.8** | 30.2 | **93.5** |
| BLAST (genome) | 64.0 | 44.9 | 36.5 | 91.5 |

Average prediction over all reads from a genome are used as a prediction for a given species. BLAST (reads) corresponds to a majority vote over all reads from a genome, and BLAST (genome) to the majority vote over all contigs. The negative class is more numerous in the dataset, which affects the precision values. The retuned classifier uses the threshold of 0.45 instead of the default 0.5 and achieves the best balanced accuracy. BAcc. – balanced accuracy, Prec. – precision; Rec. – recall, Spec. – specificity.

### Supplementary Note 4: Relationship between $s(E[z])$ and $E[s(z)]$

#### Methods

Averaging the activations of the penultimate layer for a set of reads belonging to the same genome and passing the output to the final layer, we obtain the value $s(E[z])$, where $s(x) = 1/(1 + e^{-x})$ is the sigmoid activation and $z = \mathbf{w}^\top \mathbf{h} + b$, for the penultimate activation vector $\mathbf{h}$, weights of the last layer $\mathbf{w}$ and its bias $b$. Note that this is *not* mathematically equivalent to $E[s(z)]$, or the expected output of the network for reads originating from a given genome, which was used for the genome-level predictions in previously published literature [Bartoszewicz et al., 2020, 2021b], as well as in this study. However, we suspected that although usually $s(E[z])$ and $E[s(z)]$ are not identical, those terms could have similar values in practice, as they both express a notion of aggregating information contained in individual reads to yield a prediction for a full genome. If this is the case, mean activations for a set of reads originating from the same genome can be used as if they were the internal genome representations of our species-level classifiers, even though the models rely on aggregating read-level predictions. To investigate the relationship between $s(E[z])$ and $E[s(z)]$, we performed additional experiments.

We speculated that average activations of the penultimate layer $E[\mathbf{h}]$ could be used as representations of full genomes, even though our genome-level classifiers do not explicitly use $E[\mathbf{h}]$, but rely on average per-read predictions $E[s(z)]$. Note that $E[s(z)]$ represents averaging in in the space of predicted probabilities ('proba'-average), while $s(E[z])$ is averaging in the logit space ('logit'-average). To check if $s(E[z]) \approx E[s(z)]$, we plotted the 'logit'-average predictions for each species against their 'proba'-average equivalents. As any effects found could be dataset-dependent, we perform this not only on the fungal validation and test sets, but the 'novel viruses' dataset used for the DeePaC-vir networks (and consequently, also for our multi-class classifiers). Further, we use simulated logit values ($z$) to gain deeper insight into the relationship between both versions of read averaging. To accurately model the problem of aggregating predictions for individual species in the context of binary classifications, we first defined two classes, '0' and '1'. We then simulated 100 'species' belonging to each class. Each species $S$ is described by a distribution of logit values $z \sim \mathcal{N}(\mu_S, \sigma_S^2)$, where $\mu_S$ and $\sigma_S$ are the mean and standard deviation of $z$ for that particular species and $\mathcal{N}$ is the normal distribution. To generate $\mu_S$ and $\sigma_S$ for each species, we sample from additional distributions: $\mu_S \sim \mathcal{N}(\mu_c, \sigma_c^2)$ and $\sigma_S \sim HalfNormal(\mu_\sigma, \sigma_\sigma^2)$, where $HalfNormal$ is the half-normal distribution and $c$ is the class index for classes '0' and '1'. Note that for simplicity, we sample $\sigma_S$ from the same distribution irrespective of the species's class. Finally we generate 1000 values of $z$ per species (corresponding to 1000 reads) and plot the 'logit'-averages $s(E[z])]$ against the 'proba'-averages $E[s(z)]$. Hence, to each species belonging to a given class, we assign its own mean and standard deviation used to generate read representations according to the species's class. First, we perform two experiments using the following parameters:

- $\mu_0 = -3, \mu_1 = 3, \sigma_0 = \sigma_1 = 2, \mu_\sigma = 0.25, \sigma_\sigma = 0.25$
- $\mu_0 = -3, \mu_1 = 3, \sigma_0 = \sigma_1 = 2, \mu_\sigma = 3, \sigma_\sigma = 1$

This models two well-separated classes, and the only difference between the two settings is the distribution of within-species variances of $z$.

We then test if the relationship between 'logit'- and 'proba'-average changes if no separable classes are present in the dataset. To this end, we perform two additional experiments with only one 'class', '0':

- $\mu_0 = 0, \sigma_0 = 3, \mu_\sigma = 0.25, \sigma_\sigma = 0.25$
- $\mu_0 = 0, \sigma_0 = 3, \mu_\sigma = 3, \sigma_\sigma = 1$

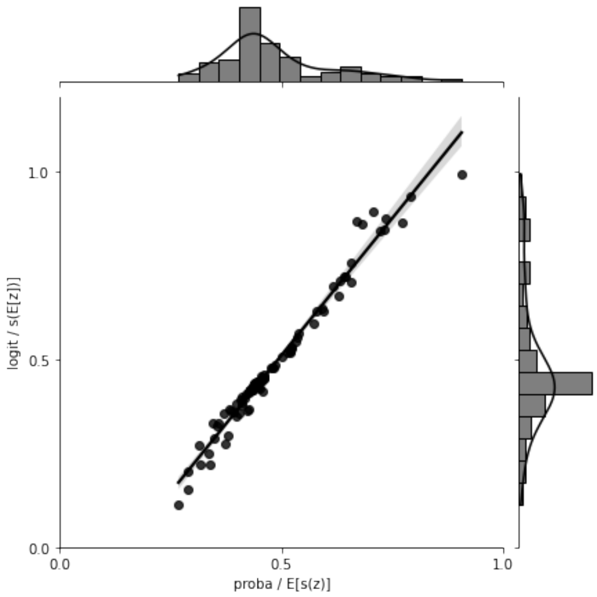


(a) Fungal validation dataset

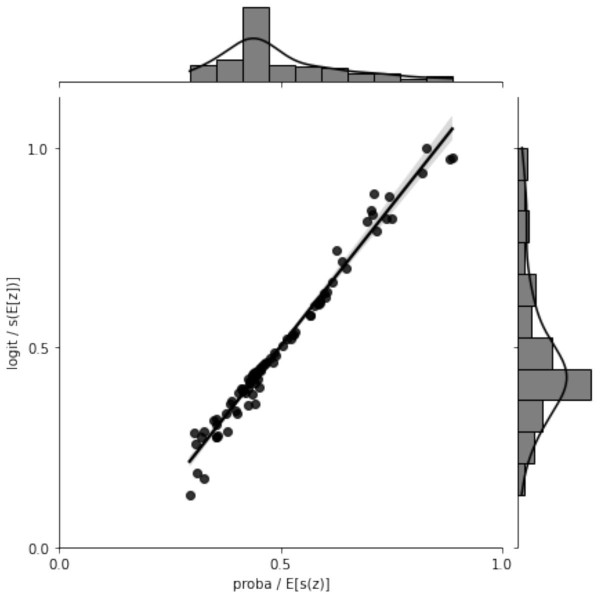


(b) Fungal test dataset

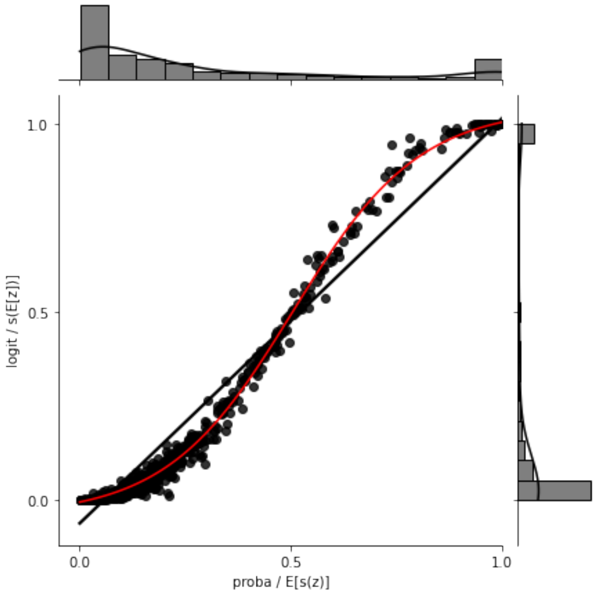


(c) 'Novel viruses' test dataset

Figure S2: Comparison of relationships between 'logit'-average ($s(E[z])$ and 'proba'-average ($E[s(z)]$) predictions. Real data.

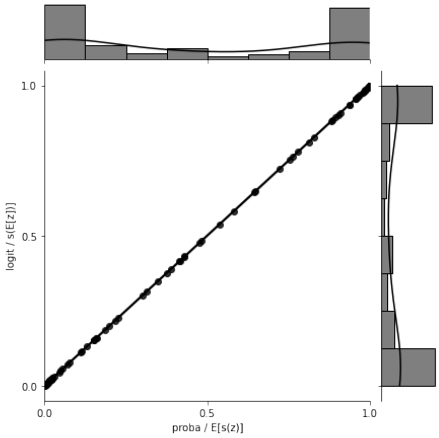


(a) Two classes, well separated, low within-species variances

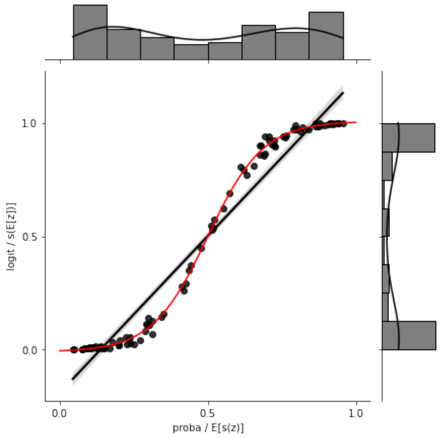


(b) Two classes, well separated, high within-species variances

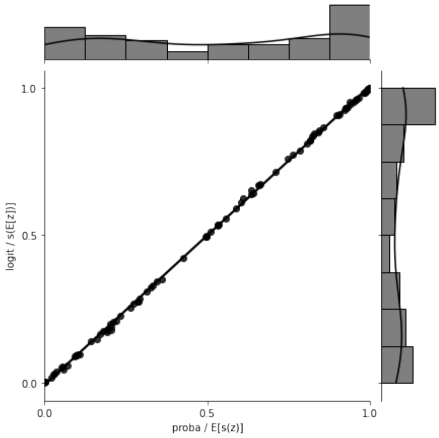


(c) No class distinction, low within-species variances

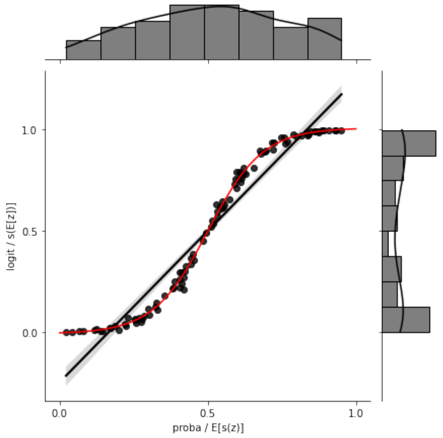


(d) No class distinction, high within-species variances

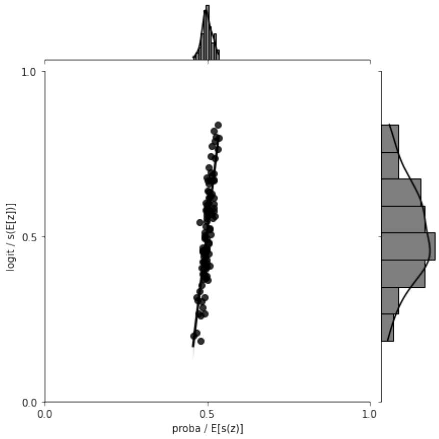


(e) No species signal, extremely high per-read variance

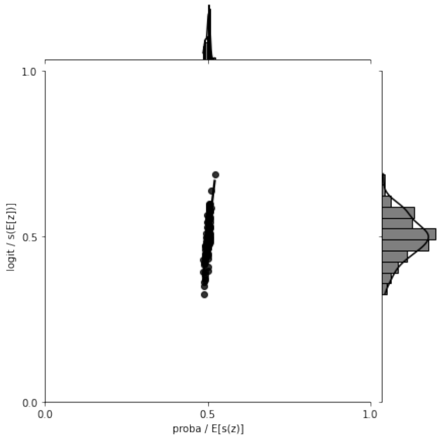


(f) No species signal, extremely high per-read variance, increased coverage

Figure S3: Comparison of relationships between 'logit'-average ($s(E[z])$ and 'proba'-average ($E[s(z)]$) predictions. Simulated $z$ values.

### Supplementary Note 5: Fungal pathogenic potential prediction

Table S3: Classification performance comparison between BLAST and the ResNet model on single reads, read pairs and genomes from the test dataset.

| | | BAcc. | Prec. | Rec. | Spec. | AUC | AUPR | Pred. |
|---|---|---|---|---|---|---|---|---|
| Single reads | ResNet | 64.7 | 65.8 | **61.3** | 68.1 | 70.9 | 72.0 | **100.0** |
| | BLAST | **66.2** | **92.6** | 61.1 | **71.3** | - | - | 77.6 |
| Read pairs | ResNet | 68.5 | 69.8 | **65.1** | 71.9 | 75.7 | 76.5 | **100.0** |
| | BLAST | **69.7** | **94.5** | 62.3 | **77.2** | - | - | 79.8 |
| Genomes | ResNet | 88.4 | 77.5 | **91.2** | 85.7 | 95.1 | 90.1 | **100.0** |
| | BLAST (reads) | **90.3** | **90.6** | 85.3 | **95.2** | - | - | **100.0** |
| | BLAST (contigs) | 89.5 | 87.8 | 85.3 | 93.7 | - | - | **100.0** |

| | | Time (CPU) | Time (GPU) |
|---|---|---|---|
| Single reads | ResNet | **25 min.** | **3 min.** |
| | BLAST | 181 min. | - |
| Read pairs | ResNet | **50 min.** | **6 min.** |
| | BLAST | 362 min. | - |
| Genomes | ResNet | **25–50 min.** | **3–6 min.** |
| | BLAST (reads) | 181–362 min. | - |
| | BLAST (contigs) | 2190 min. | - |

The best performance value for each performance metric (row) and setting is displayed in bold. Read datasets are balanced, and the differences in performance for the first and second mate were negligible for all metrics. In genome datasets, negative class is the majority class. We use BLAST as shown in [Bartoszewicz et al., 2020]; since this only returns binary labels, AUC and AUPR are not defined. Overall, BLAST is much more precise, but the ResNet yields only slightly less accurate predictions for all reads, even those impossible to match with BLAST, in a fraction of the time. We comment more on this fact in Supplementary Note 5. AUC and AUPR are marginally higher (95.2 and 90.2) for the genome-level ResNet if only first mates are used compared to using both mates (95.1 and 90.1). Other metrics are equal, and the computation time is 50% lower. BAcc. – balanced accuracy (equivalent to accuracy on read sets), Prec. – precision; Rec. – recall, Spec. – specificity, AUC – area under the ROC curve, AUPR – area under the PR curve, Pred. – prediction rate; Time (CPU) – prediction time on 2x AMD EPYC 7742 (256 threads); Time (GPU) – prediction time on 1x Tesla V100 GPU (not possible for BLAST).

Table S4: Classification performance comparison between BLAST and the ResNet model on single reads and read pairs from the temporal dataset.

| | | BAcc. | Prec. | Rec. | Spec. | AUC | AUPR | Pred. |
|---|---|---|---|---|---|---|---|---|
| Single reads | ResNet | 68.9 | 69.6 | 66.9 | **70.8** | 74.9 | 74.6 | **100.0** |
| | BLAST | **71.2** | **87.4** | **81.3** | 61.2 | - | - | 82.6 |
| Read pairs | ResNet | **74.2** | 74.6 | 73.4 | **75.1** | 81.8 | 82.1 | **100.0** |
| | BLAST | 71.9 | **87.4** | **77.5** | 66.3 | - | - | 80.1 |

The best performance value for each performance metric (row) and setting is displayed in bold. While the read number per class is balanced, the dataset is small and extremely imbalanced on the genome level – only a single human pathogen genome and 14 non-human pathogen genomes were added to GenBank in the time window of our temporal benchmark. Therefore, the presented results are not fully representative, and the higher performance of the ResNet on read pairs cannot be guaranteed in general. However, the overall results are similar to those obtained for our main held-out test dataset. BAcc. – balanced accuracy (equivalent to accuracy on read sets), Prec. – precision; Rec. – recall, Spec. – specificity, AUC – area under the ROC curve, AUPR – area under the PR curve, Pred. – prediction rate.

Table S5: Effects of integrating multiple classes on the classification performance on non-fungal datasets, read pairs.

| | | Acc. | Prec. | Rec. | Spec. | AUPR |
|---|---|---|---|---|---|---|
| Bacteria | 4-class ensemble | 85.1 | **84.1** | 87.2 | **83.0** | 88.3 |
| | 4-class | 83.6 | 80.9 | 89.0 | 78.1 | 88.6 |
| | 3-class | 84.0 | 80.6 | 89.7 | 78.2 | **90.0** |
| | 2-class [Bartoszewicz et al., 2021a] | **87.3** | 83.6 | **92.7** | 81.8 | 89.0 |
| | BLAST | 70.0 | **84.1** | 86.6 | 53.5 | - |
| Viruses | 4-class ensemble | 81.4 | 94.3 | 90.9 | 71.8 | **98.1** |
| | 4-class | 79.0 | 93.4 | **91.2** | 66.7 | 97.9 |
| | 3-class | 81.9 | 93.3 | 91.1 | 72.7 | 97.8 |
| | 4-class ensemble, reassigned | 89.6 | 95.3 | 89.3 | 90.0 | **98.1** |
| | 4-class, reassigned | 89.6 | 94.7 | 89.6 | 89.7 | 97.9 |
| | 3-class, reassigned | **91.8** | 95.1 | 88.1 | **95.4** | 97.8 |
| | 2-class [Bartoszewicz et al., 2021a] | 90.3 | 94.7 | 85.5 | 95.2 | 97.2 |
| | BLAST | 80.6 | **98.4** | 79.1 | 82.2 | - |

The 4-class classifier includes the human-pathogenic fungi class along the three viral and bacterial classes included in the 3-class classifier. The 2-class classifiers are the original DeePaC ResNets [Bartoszewicz et al., 2021a] for bacteria and viruses accordingly. We use BLAST as shown in [Bartoszewicz et al., 2020, 2021b]; since this only returns binary labels, AUPR is not defined. The multi-class models do not assume a purely bacterial or viral sample, so they are more flexible than the 2-class networks – a single classifier can be used for both datasets without retraining, although a minor cost in performance should be expected of a more general approach. For the viral dataset, assignments to the bacterial pathogens class may be assumed to actually reflect bacteriophages infecting those bacteria due to high bacteriophage-host sequence similarity. Classifiers reassigning the bacterial assignments into non-pathogen class based on this assumption are listed as 'reassigned'.

The multi-class models and the specialized binary classifiers perform comparably well, even though the latter can only be used in very specific situations. Integrating fungi into the classifiers does not significantly impact the overall accuracy. The main challenge in multi-class pathogenic potential prediction seems to be actually differentiating between bacteriophages and the pathogenic bacteria that they infect. The multi-class networks confuse 18–21% of viral non-pathogen reads for pathogenic bacteria. However, their accuracy is still similar to BLAST's, even though BLAST was used with a purely viral reference database, and hence would always classify ambiguous bacterial or viral sequences as viral. This problem is unrelated to the fungal database presented here, and can be circumvented by assuming that spurious assignments to bacteria should be reassigned to bacteriophages in the case of a purely viral dataset. The resulting networks achieve accuracy similar to that of the simple binary classifiers, while still being capable of multi-class predictions for more complex datasets. They also outperform BLAST, which represents the estimated upper limit on the accuracy of pathogen detection by finding the closest taxonomic matches, by a wide margin.

Acc. – accuracy, Prec. – precision; Rec. – recall, Spec. – specificity, AUPR – area under the PR curve (calculated for the respective positive class).

Table S6: Effects of integrating multiple classes on the classification performance on the multi-class dataset, read pairs.

| | | Acc. | F1 | Prec. | Rec. | AUPR |
|---|---|---|---|---|---|---|
| All classes | 4-class ensemble | 87.6 | 87.7 | 87.7 | 87.6 | 93.4 |
| | 4-class | 86.6 | 86.7 | 86.8 | 86.6 | 92.8 |
| | 3-class | 85.5 | 85.6 | 85.8 | 85.5 | 92.4 |
| Non-pathogens | 4-class ensemble | **77.4** | **78.7** | 80.1 | **77.4** | 86.7 |
| | 4-class | 72.5 | 76.7 | 81.5 | 72.5 | 85.3 |
| | 3-class | 75.5 | 78.6 | **81.8** | 75.5 | **87.6** |
| Path. bacteria | 4-class ensemble | 87.2 | **85.1** | **83.2** | 87.2 | 90.4 |
| | 4-class | 89.0 | 83.8 | 79.1 | 89.0 | 90.4 |
| | 3-class | **89.7** | 84.2 | 79.3 | **89.7** | **91.1** |
| Human viruses | 4-class ensemble | 90.9 | **93.7** | **96.7** | 90.9 | 98.4 |
| | 4-class | **91.2** | 93.4 | 95.7 | **91.2** | 98.1 |
| | 3-class | 91.1 | 93.6 | 96.3 | 91.1 | **98.5** |
| Fungi | 4-class ensemble | **95.0** | **92.9** | **90.9** | **95.0** | **97.9** |
| | 4-class | 93.7 | 92.2 | 90.7 | 93.7 | 97.4 |

The 4-class classifier includes the fungi class along the three viral and bacterial classes included in the 3-class classifier. The best performance for each class is marked in bold. In this setting, the true positive rate corresponds to the rate of correct assignments within a given class. Hence, recall is equal to accuracy for each individual class. We use the F1 score as an additional measure. Integrating fungi into the classifiers does significantly impact the overall accuracy for any of the non-fungal classes, and the 4-class ensemble offers a more balanced performance on putative non-pathogen reads than the single-network alternative. The metrics labelled as 'All classes' were calculated for all four classes for the 4-class models, and for the three relevant classes for the 3-class model. Acc. – accuracy, F1 – F1 score, Prec. – precision, Rec. – recall, AUPR – area under the PR curve.

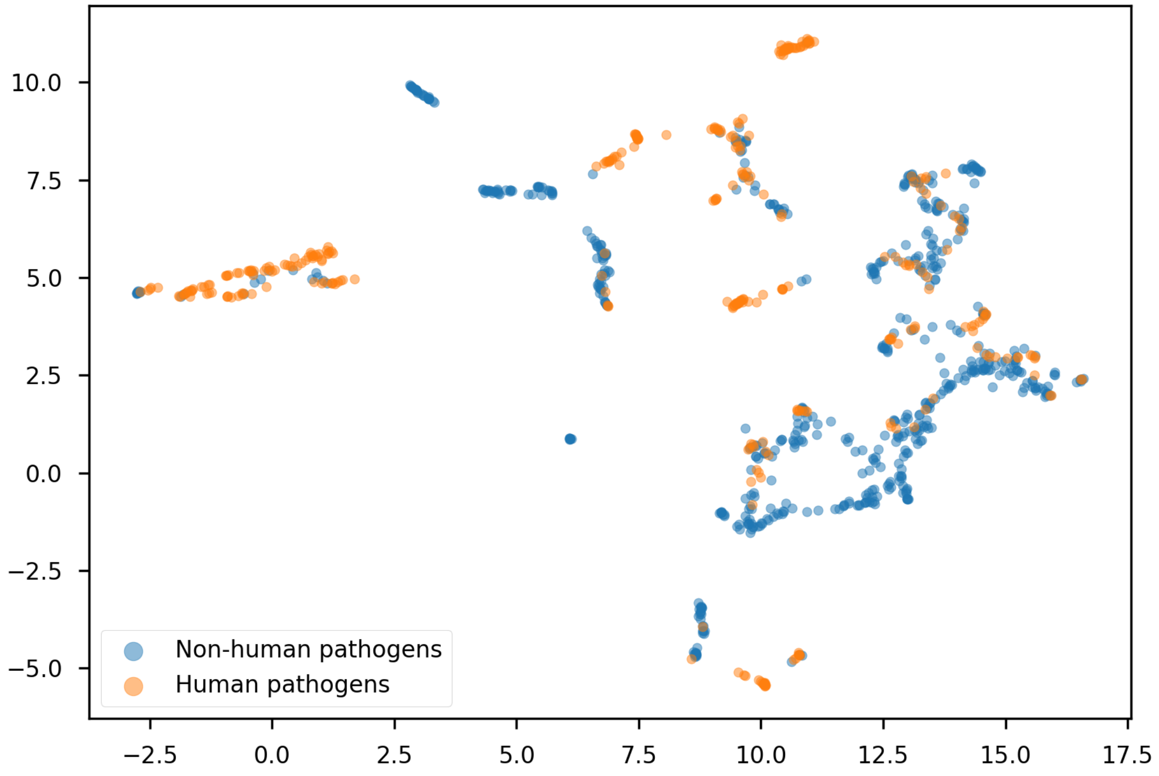


Figure S4: UMAP embeddings of the learned genome representations for the core database; true labels.

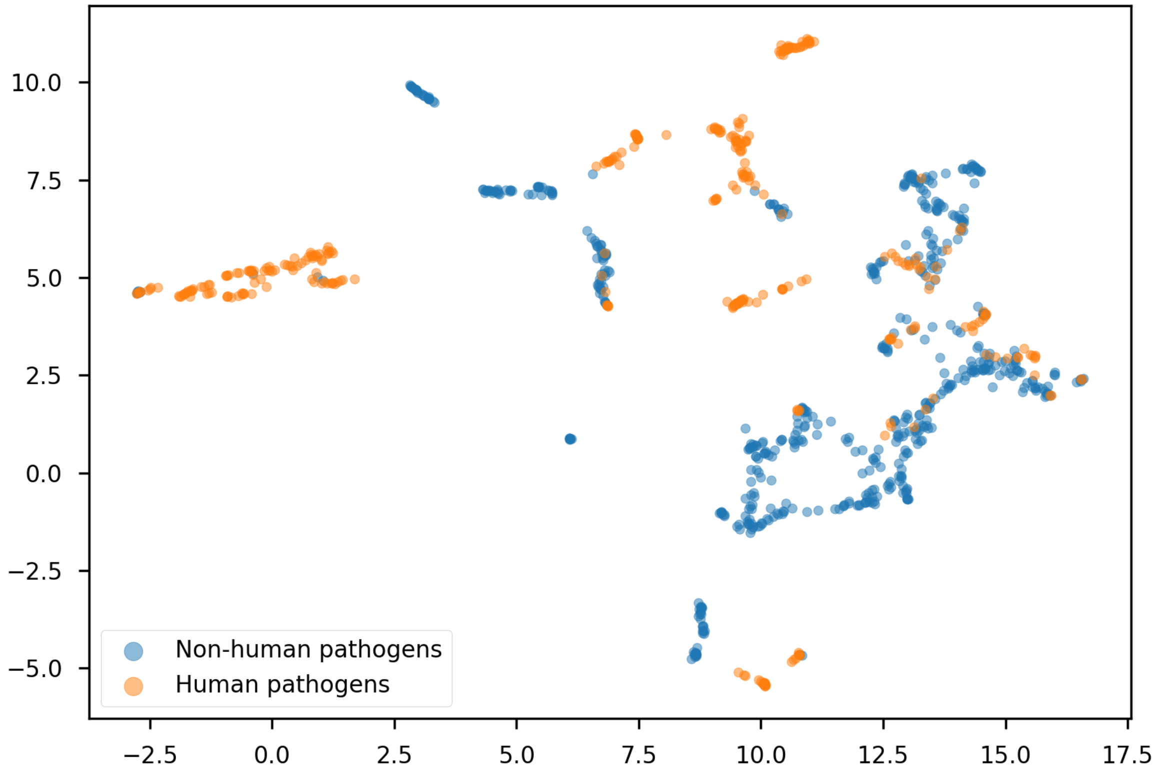


Figure S5: UMAP embeddings of the learned genome representations for the core database; labels predicted by the ResNet (retuned threshold). Most 'positive' members of otherwise 'negative' clusters are correctly detected.

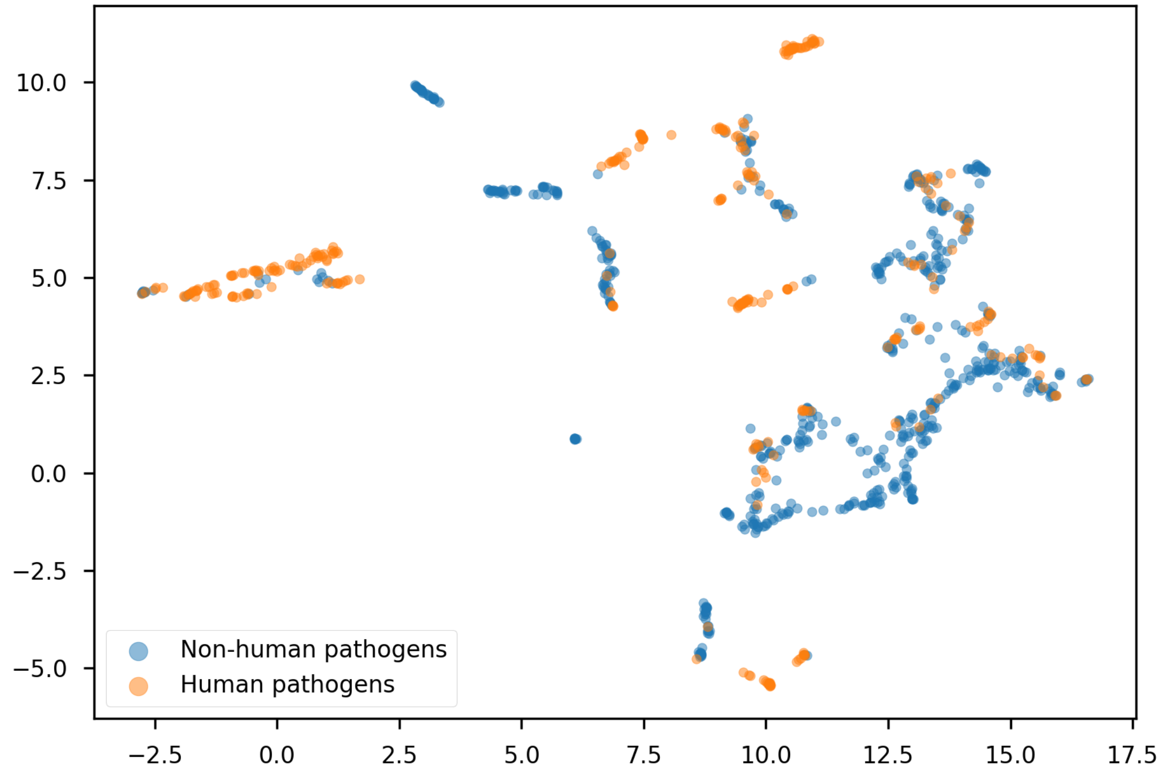


Figure S6: UMAP embeddings of the learned genome representations for the core database; labels predicted by BLAST. Training species are present in the reference database, so predicting labels for them is easy (99.9% accuracy).

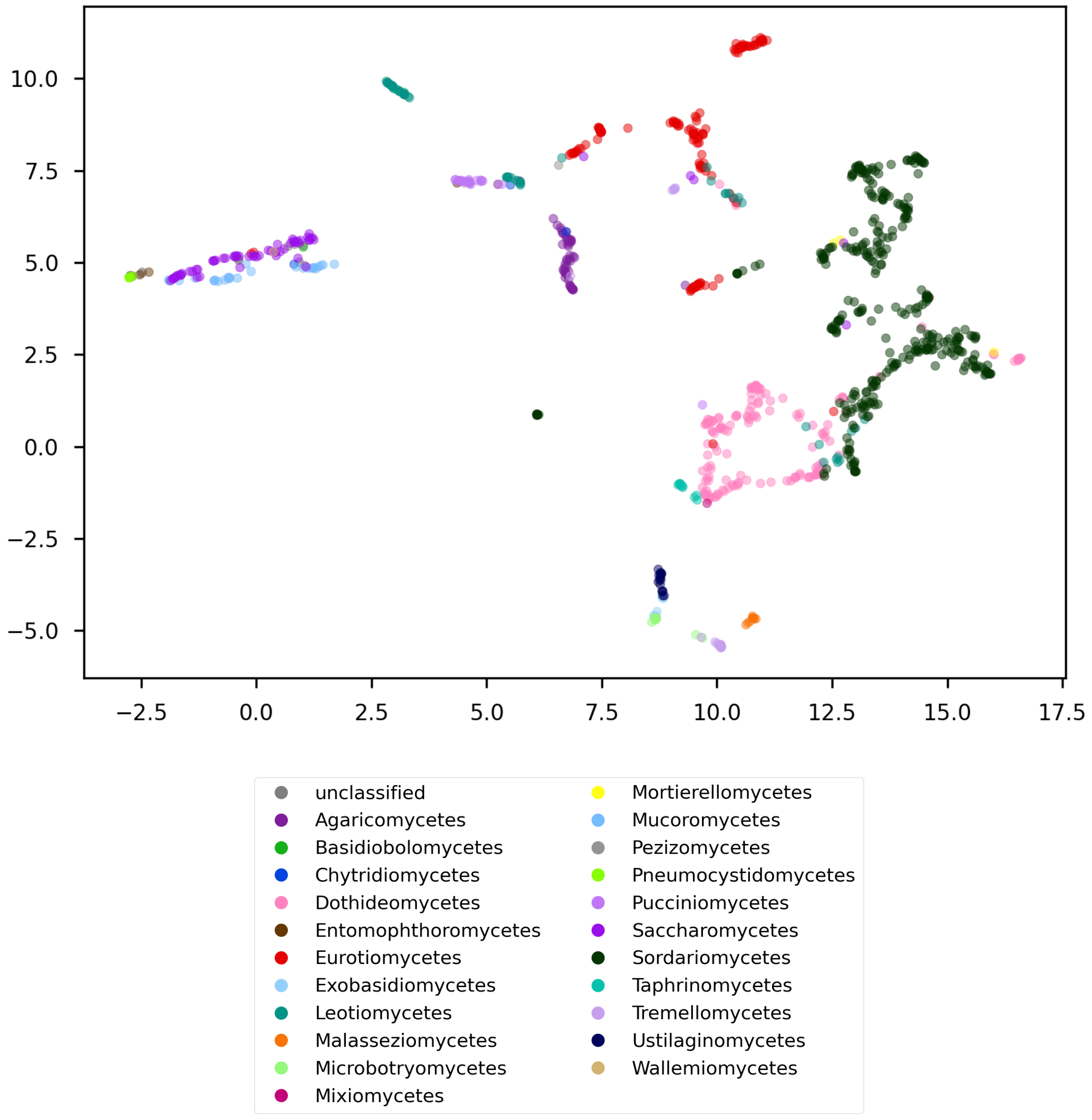


Figure S7: UMAP embeddings of the learned genome representations for the core database; taxonomic rank: class. In general, related species are close to each other in this space, but some taxonomic units are distributed among more than one cluster (e.g. *Eurotiomycetes*, in red), and some clusters contain members of distant taxa (e.g. the leftmost cluster).

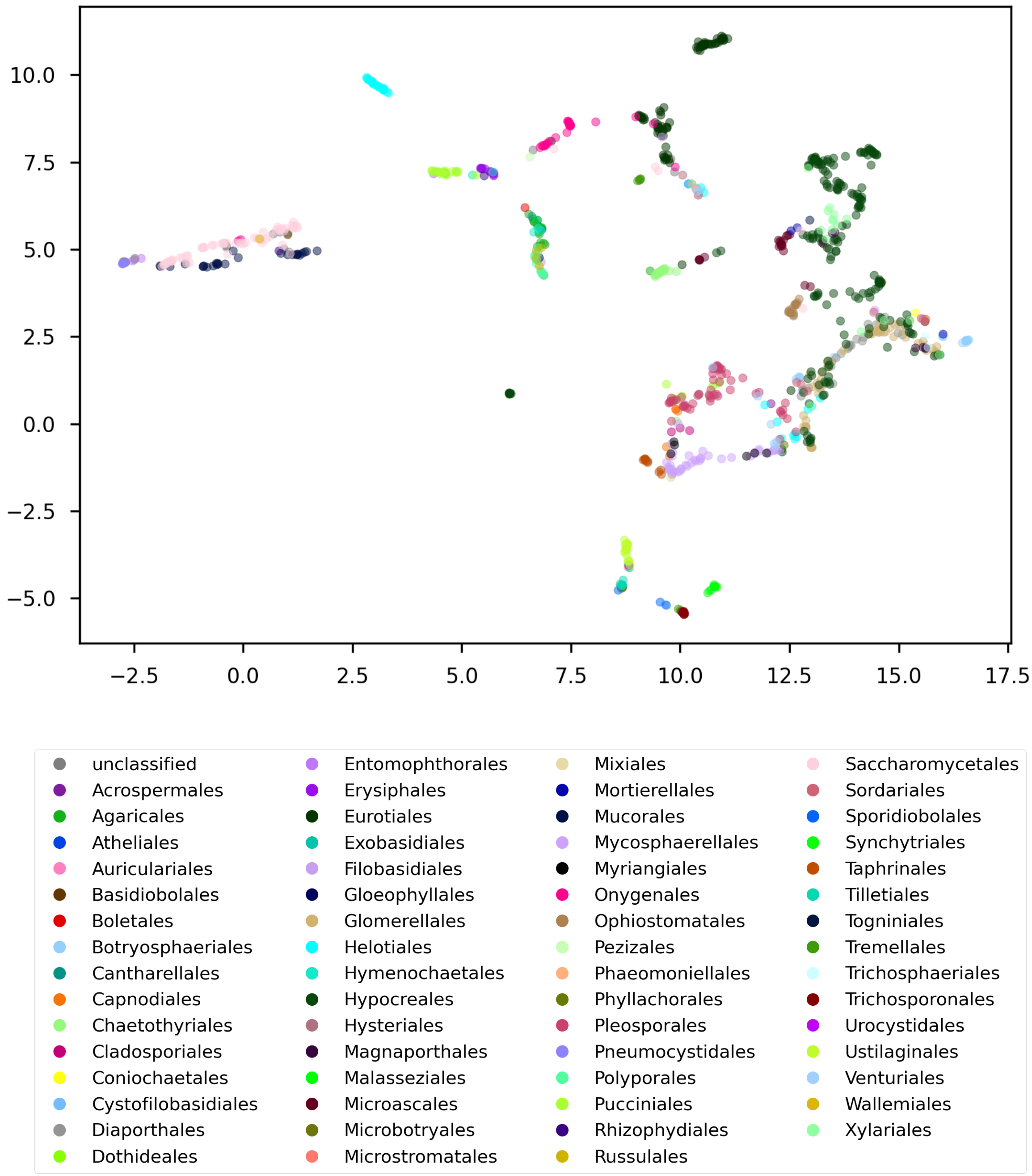


Figure S8: UMAP embeddings of the learned genome representations for the core database; taxonomic rank: order.

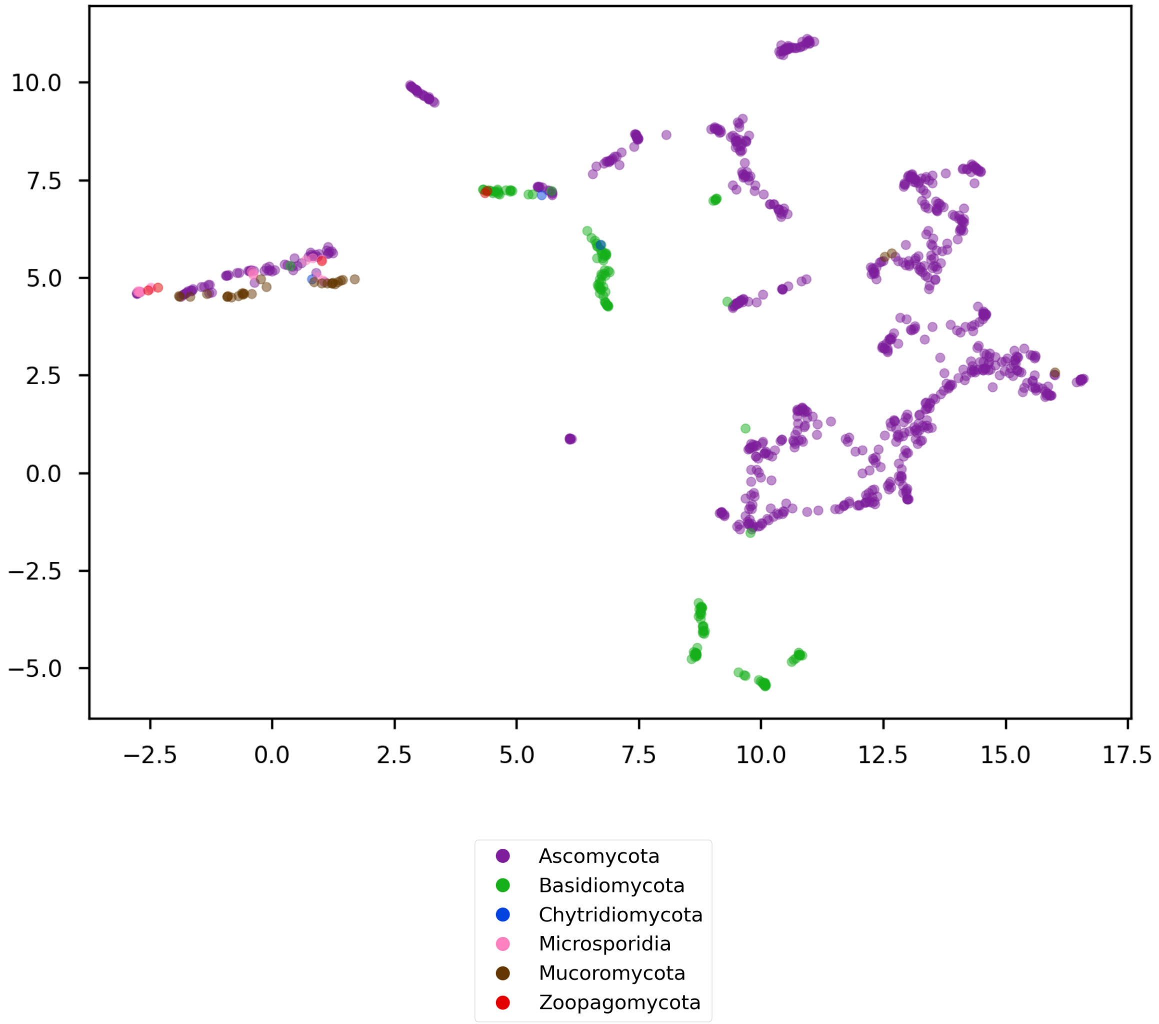


Figure S9: UMAP embeddings of the learned genome representations for the core database; taxonomic rank: phylum.

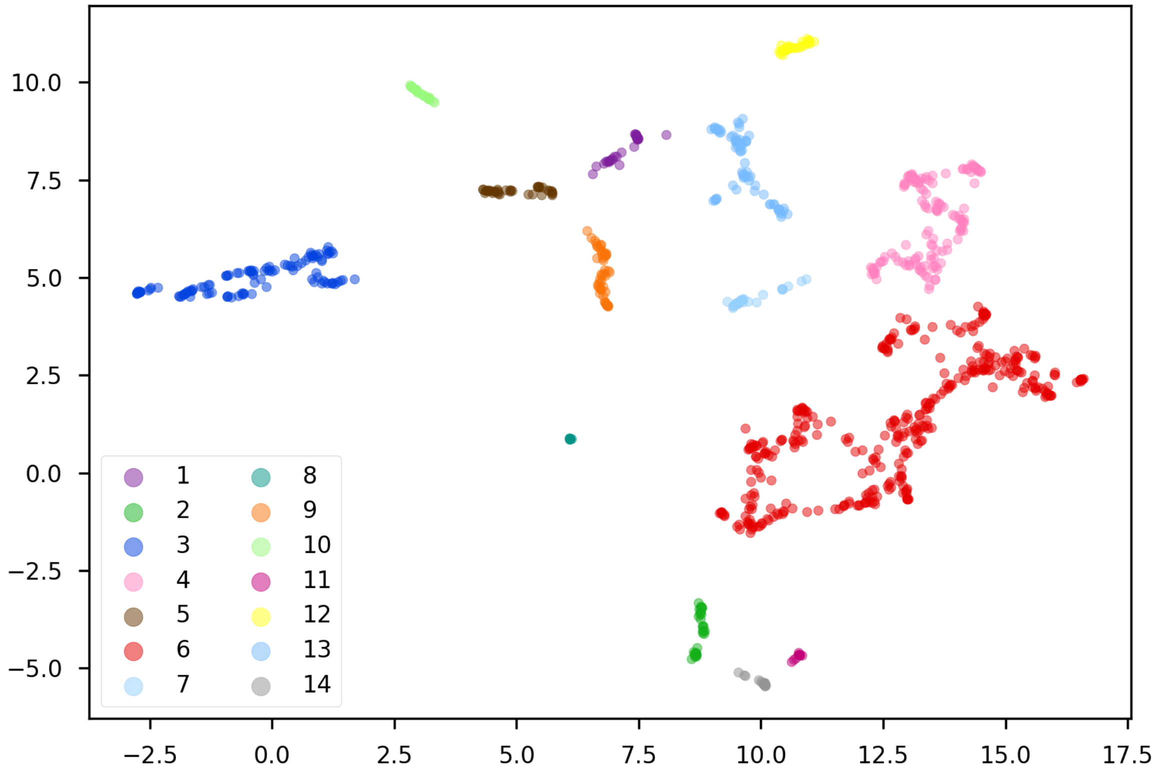


Figure S10: UMAP embeddings of the learned genome representations for the core database. Assignment to clusters detected with single-linkage agglomerative clustering. Clusters numbered arbitrarily.

Table S7: *C. auris* sequencing run, SRA accession SRR17577041. Binary classifiers (host group prediction).

| | | Acc. | Rec. | Pred. |
|---|---|---|---|---|
| First read | 2-class (ours) | **77.4** | **77.4** | **100.0** |
| | BLAST | 46.9 | 46.9 | 52.0 |
| | STAT | 0.1 | 0.1 | 7.7 |
| Both reads | 2-class (ours) | **78.4** | **78.4** | **100.0** |
| | BLAST | 50.1 | 50.1 | 56.9 |
| | STAT | 0.1 | 0.1 | 5.4 |

As this is a pure pathogen sample, accuracy and recall are equivalent. We report performance metrics for the first mate and both mates. The mean sequencing quality for the second mate was low (below 28). This is a problem especially for STAT, which performs worse if both reads are counted than if only the first, higher-quality mate is considered. BLAST and ResNets are more robust. ResNets are the best methods overall. Interestingly, 2-class ResNet performance for this species is above average (see Table S3). Perhaps this is because *Candida* species are placed in a cluster of conserved pathogens (see Supp. Note 6). Acc. – accuracy, Rec. – recall, Pred. – prediction rate (a fraction of reads with any hits).

#### Supplementary Note 7: Performance on real data

Obtaining ground truth for genuine metagenomic samples is very challenging without spiking them with known organisms or corresponding reads. As our test sets consist of reads simulated from a mixture of different species (and viruses), they effectively function as mock metagenomic samples. Simulated reads are commonly used to evaluate computational approaches for read classification (e.g. [Wood et al., 2019, Katz et al., 2021]. To show that the methods discussed here work also on real data, we focus on a single-species sequencing run, where ground truth is available. We selected a *C. auris* sequencing run (SRA accession SRR17577041) and analyzed it in two scenarios: a) assuming it is a novel fungal species, predict whether it could infect humans (binary classification, as in Table S3-Table S4) b) assuming it is an unknown agent originating from a patient, predict its correct pathogen group (multi-class setting, as in Table 1 and Table S6). In addition to the corresponding ResNets and BLAST databases, we also used STAT [Katz et al., 2021], a k-mer tool used by the NCBI to generate taxonomy reports for all SRA archives.

To evaluate STAT, we downloaded its reference database published along the paper and run a *C. auris* strain exclusion study as described by the authors. In the host prediction setting Table S7, we counted all STAT species-level hits to any pathogenic fungal species as positives, but in the multi-class pathogen detection task (Table 2), we assumed that all reads classified as fungi can be counted as successful detection. As STAT has no way of jointly classifying read pairs, we treated both mates as independent reads also when comparing to BLAST and ResNet predictions for read pairs. Note that in contrast to our BLAST evalutation, STAT reference database contained 248,426 TaxIds (see Katz et al. [2021]), and only *C. auris* was effectively removed from it for the strain exclusion study.
